# Purification, characterization and influence on membrane properties of the plant-specific sphingolipids GIPC

**DOI:** 10.1101/2020.10.01.313304

**Authors:** Adiilah Mamode Cassim, Yotam Navon, Yu Gao, Marion Decossas, Laetitia Fouillen, Axelle Grélard, Minoru Nagano, Olivier Lambert, Delphine Bahammou, Pierre Van Delft, Lilly Maneta-Peyret, Françoise Simon-Plas, Laurent Heux, Giovanna Fragneto, Jenny C. Mortimer, Magali Deleu, Laurence Lins, Sébastien Mongrand

## Abstract

The plant plasma membrane (PM) is an essential barrier between the cell and the external environment. The PM is crucial for signal perception and transmission. It consists of an asymmetrical lipid bilayer made up of three different lipid classes: sphingolipids, sterols and phospholipids. The most abundant sphingolipids in the plant PM are the Glycosyl Inositol Phosphoryl Ceramides (GIPCs), representing up to 40% of total sphingolipids, assumed to be almost exclusively in the outer leaflet of the PM. In this study, we investigated the structure of GIPCs and their role in membrane organization. Since GIPCs are not commercially available, we developed a protocol to extract and isolate GIPC-enriched fractions from eudicots (cauliflower and tobacco) and monocots (leek and rice). Lipidomic analysis confirmed the presence of different long chain bases and fatty acids. The glycan head groups of the different GIPC series from monocots and dicots were analysed by GC-MS showing different sugar moieties. Multiple biophysics tools namely Langmuir monolayer, ζ-Potential, light scattering, neutron reflectivity, solid state ^2^H-NMR and molecular modelling were used to investigate the physical properties of the GIPCs, as well as their interaction with free and conjugated phytosterols. We showed that GIPCs increase the thickness and electronegativity of model membranes, interact differentially with the phytosterols species and regulate the gel-to-fluid phase transition during temperature variations.

## Introduction

The plant plasma membrane (PM) contains three main classes of lipids: phytosterols, sphingolipids and phospholipids, all with a high level of molecular complexity, see (Cacas et al., 2016) (Yetukuri et al, 2008). In plants, the major sphingolipid subclass is the Glycosyl Inositol Phosphoryl Ceramides (GIPCs). GIPCs were discovered in plants and fungi during the 1950’s (Carter et al., 1958). The structural diversity of GIPCs lies in the hydroxylation, degree and position of saturation of their fatty acid (FA) chain and long chain base (LCB) and glycosylation (Pata et al., 2010). Plant GIPCs predominantly consist of a t18:0 or t18:1 LCB (trihydroxylated saturated or monounsaturated) amidified to a Very Long Chain Fatty Acid (VLCFA) or 2-hydroxylated VLCFA (hVLCFA) to form a ceramide (Cacas et al., 2016)(Buré et al., 2011).

The GIPC head group linked to the ceramide consists of a phosphate bound to an inositol, forming the inositol phosphoryl ceramide (IPC) backbone, which is then further substituted with further sugar moieties. A broad study of the GIPC polar heads of 23 plant species from algae to monocots showed that polar head structures are largely unknown and vary widely across different biological taxa (Cacas et al., 2013). GIPCs are classified into series, based on the degree of glycosylation of their polar head group (Buré et al., 2011). In plants, all GIPCs characterized to date have a glucuronic acid (GlcA) as the first sugar on the IPC, followed by at least one more sugar unit of varying identity. For example, GIPC series A is defined as one monosaccharide addition to the GlcA-IPC form (Buré et al., 2011). In the 1960s, the first characterization of a GIPC structure from *Nicotiania tabacum* (tobacco) was described (Carter et al., 1958)(Carter & Kisic, 1969) (Carter & Koob, 1969) (Hsieh, Lester, & Laine, 1981) (Kaul & Lester, 1978). The GIPC extraction method required hundreds of kilograms of plant material and litres of solvents. They fully resolved the exact number and type of sugars as well as the nature of the sugar bond. For instance, the reported series A GIPC still has the best described structure to date: GlcNAc(α1➔4)GlcA(α1➔2)inositol-1-O-phosphorylceramide, see Figure 1A. Additional sugar moieties were described, such as glucosamine (GlcN), *N*-acetyl-glucosamine (GlcNAc), arabinose (Ara), galactose (Gal) and mannose (Man), which may lead to observed glycan patterns of three to seven sugars, the so-called GIPC series B to F. It is noteworthy that Kaul and Lester calculated the ratio between carbohydrate/LCB/inositol in purified polyglycosylated GIPCs and showed that they may contain up to 19–20 sugars (Kaul & Lester, 1975), which opens a very large field of investigation. Polyglycosylated GIPCs found in *Zea mays* (corn) seeds and *Erodium* displays branched polar heads (Sperling & Heinz, 2003)(Buré et al., 2016). GIPC series are species- and tissue-specific. In Arabidopsis, the GIPC series A headgroup Man-GlcA-IPC is predominant in leaves and callus (Mortimer et al., 2013) (Fang et al., 2016), whereas a complex array of *N*-acetyl-glycosylated with up to three pentose units are present in pollen (Luttgeharm et al., 2015). Amino- and *N*-acylated-GIPCs are found in Arabidopsis seeds and oil (Tellier et al., 2014). GlcN(Ac)-GlcA-IPC are mainly found in rice and tobacco (Buré et al., 2011) (Nagano et al., 2016). In monocots, the predominant GIPC series is series B (Buré et al., 2011), their core structures are yet to be deciphered.

**Figure 1.**
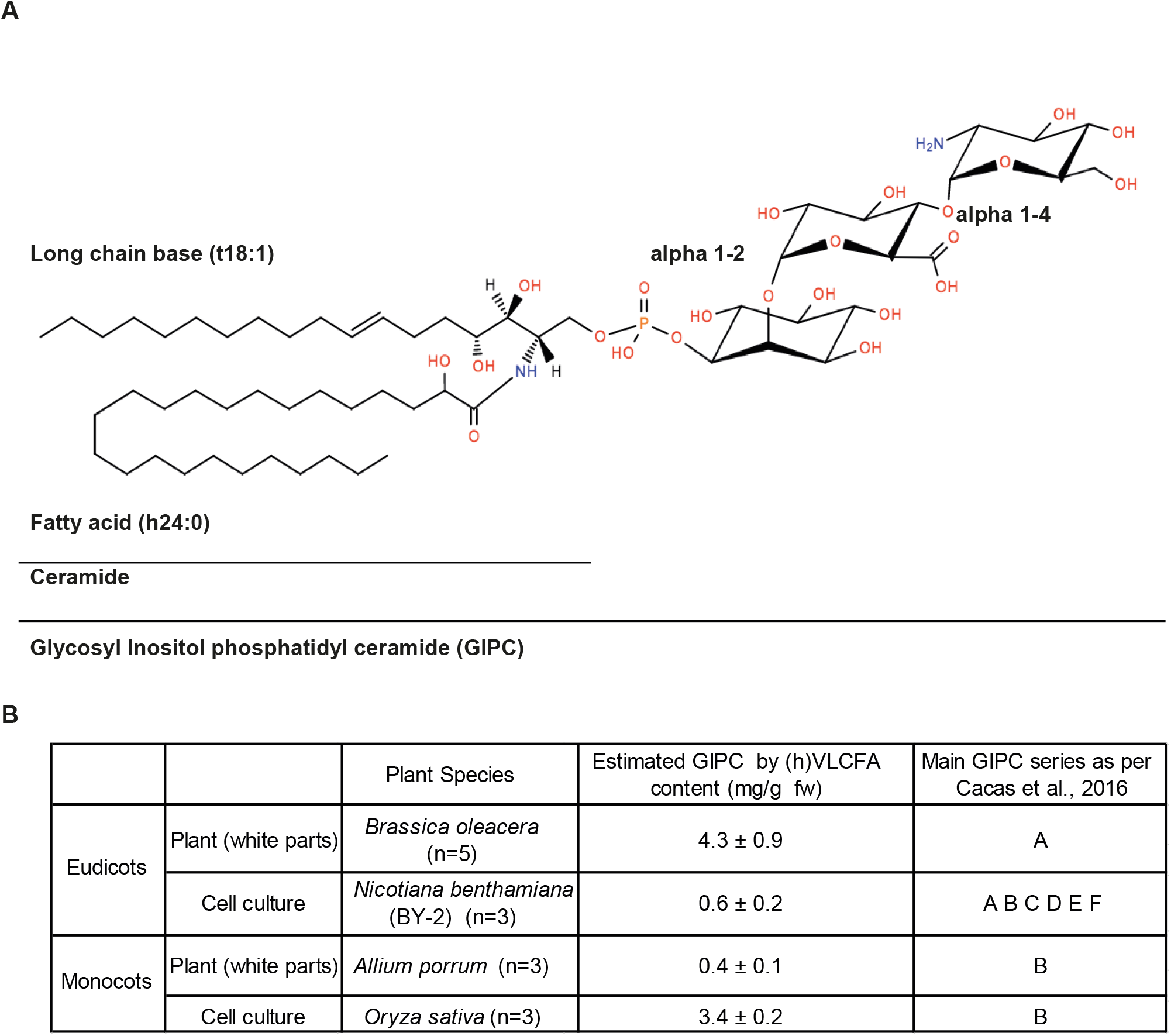
A, Structure of GIPC series A (2 sugars after the inositol group). B, GIPC content of different plant species: *Brassica oleracea* (cauliflower), *Nicotiana tabacum* (BY-2 cell culture), *Allium porrum* (leek) and *Oryza sativa* (rice cell culture). The GIPC content in mg per g of fresh weight was estimated by calculating the proportion of (h)VLCFA (hydroxylated Very Long Chain Fatty Acid) as determined by fatty acid methyl ester (GC-MS). The type of GIPC was defined by HPTLC analysis based on Cacas et al., 2016. Three to five independent samples were processed.

The GIPC’s polar head is responsible for the high polarity of the GIPC, accounting for its insolubility in traditional lipid extraction solvents, such as chloroform/methanol. Consequently, they are lost in the aqueous phase or at the interface. GIPCs, although one of the fundamental components of the plant PM model have been poorly studied, in part due to the absence of commercial preparations. Recent evidence has demonstrated that a loss of the glycosylation is lethal, (Ishikawa et al., 2018) (Rennie et al., 2014), and that misglycosylation affects both abiotic and biotic stress responses, as reviewed in (Mortimer & Scheller, 2020).

Lipids are not homogeneously distributed within the PM bilayers. The lateral partitioning observed in the PM might be due to differential phase behaviours of different lipid species due to specific interactions between their different lipid species (Sampaio et al., 2009). This was reported in model membranes, using biophysical approaches and super resolution microscopy (Levental & Veatch, 2016). Lipid domains or liquid-ordered (Lo) phases are formed from saturated phospholipids and sphingolipids in the presence of sterol, while liquid-disordered (*Ld*) phases are formed mainly from unsaturated phospholipids (Baumgart et al., 2007) (Lingwood & Simons, 2010). In Lo phases, the high degree of conformational order is imposed on the acyl tails of lipids by the rigid ring structure of cholesterol. This increases the thickness of the lipid bilayer and lipid packing, although lipids remain laterally mobile (Mannock et al., 2010). Plant sterols and sphingolipid/sterol interactions have recently been reported as important determinants of lipid partitioning and organization within the PM (Beck et al., 2007; Gerbeau-Pissot et al., 2014; Grosjean et al., 2015). The plant PM contains 20–50% sterols, depending on plant species and organ (Furt et al., 2007), harbouring a wide molecular diversity including free and conjugated species and dominated by β-sitosterol, stigmasterol, and campesterol (Moreau et al., 2018). These phytosterols play significant roles in regulating the order level of the membrane such that ternary mixtures (sterol/sphingolipid/saturated phospholipid) have less temperature sensitivity to thermal variations compared to systems mimicking the lipid rafts of animal and fungi (Beck et al., 2007). β-sitosterol and campesterol have the largest effect on lipid ordering (Beck et al., 2007; Cacas et al., 2016; Grosjean et al., 2015). Conjugated forms of β-sitosterol and stigmasterol are proposed to reinforce membrane cohesion by additional attractive van der Waals interactions with the acyl chains of sphingolipids and phospholipids (Beck et al., 2007). Using environment–sensitive probes, it was shown that various phytosterols have the ability to modulate the proportion of Lo phases and membrane heterogeneity *in vivo* as *in vitro,* with the notable exception of stigmaterol (Gerbeau-Pissot et al., 2014) (Grosjean et al., 2015). Thus, GIPCs in synergy with sterols may organize and promote large ordered domains such that both have important roles in PM sub-compartmentalization and membrane dynamics (Grosjean et al., 2015).

As mentioned previously, older published protocols used large amounts of material and solvents, which is not feasible in modern labs. More recently published protocols do not yield enough material of high enough purity for structural characterization. In this project, we devised a new protocol to obtain milligram amounts of highly enriched GIPC samples from both monocots and eudicots, suitable for use in studies of GIPC structure and its role in PM organization. Using biophysics tools such as Langmuir monolayers, molecular modelling, supported lipid bilayers, giant unilamellar vesicles (GUVs), dynamic light scattering (DLS), ζ-potential, cryo-electron microscopy (cryo-EM), solid state ^2^H-NMR and neutron reflectivity, we aim to uncover the role of GIPCs, in synergy with sterols, in the plant PM organization.

## Results

### Extraction and Purification of GIPC-enriched fractions from different plant species tissues and cell culture

To assist with purifying the milligram amount of GIPCs required for analysis, we first assessed the amount of GIPCs in different plant species and tissues. We chose species/tissues which are easily and abundantly available, and quantified the non-hydroxylated VLCFA and 2-hydroxylated hVLCFA, diagnostic of plant’s GIPC (Cacas et al., 2016). Four species were selected: 2 eudicot plants: cauliflower (*Brassica oleacera, Bo*) head and *Nicotiana tabacum* (*Nt*) cell culture Bright-Yellow 2 (BY-2) and 2 monocot plants: the white leaves of leek (*Allium porrum, Ap*) and rice (*Oryza sativa, Oz*) cell culture. The white part of plant tissues and cell cultures were used to avoid contamination by the abundant plastidial lipids and pigments. Cauliflower and rice cell culture have the highest GIPC content with an estimated 4.3 mg/ml and 3.4 mg/ml per fresh weight respectively (Figure 1B). BY2 cells and leek both had a much lower GIPC content, with a mean estimated content of 0.6 mg/ml and 0.4 mg/ml per fresh weight respectively.

GIPCs were extracted from all 4 materials to get different GIPC series. To maximize the yield, several trials were performed to test the different published protocols of GIPCs. Steps from three protocols were organized to obtain the best yield of GIPC. These protocols were from (Carter & Koob, 1969), (Markham & Jaworski, 2007) and (Kaul & Lester, 1975). Figure 2 shows the extraction and purification processes to obtain GIPC-enriched fractions of cauliflower (Bo-GIPC), tobacco BY-2 (Nt-GIPC), leek (Ap-GIPC) and rice (Os-GIPC). Some fine-tuning was done to maximize the yield such as refluxing in boiling ethanol for 20 min and using large lab-made silica column to process several hundreds of grams of material (see Material and Methods). Crude sphingolipid extracts were directly dried in silica deposited on the top of the column chromatography. The column was then washed with 4 column volumes (cv) of a mix of chloroform/methanol with increasing polarity to remove sterols, glucosylceramide and phospholipids. For the elution of GIPCs, a step gradient of chloroform/methanol/water was used (Figure 2), so that molecules of increasing polarity were eluted in the last fractions. All washes and elution fractions were collected and analysed by high-performance thin layer chromatography (HPTLC) as shown in Supplementary data 1. HPTLC was a quick and reliable way to select fractions enriched with GIPCs, because it allowed the clear separation of sterols, phospholipids, and GIPC series.

**Figure 2.**
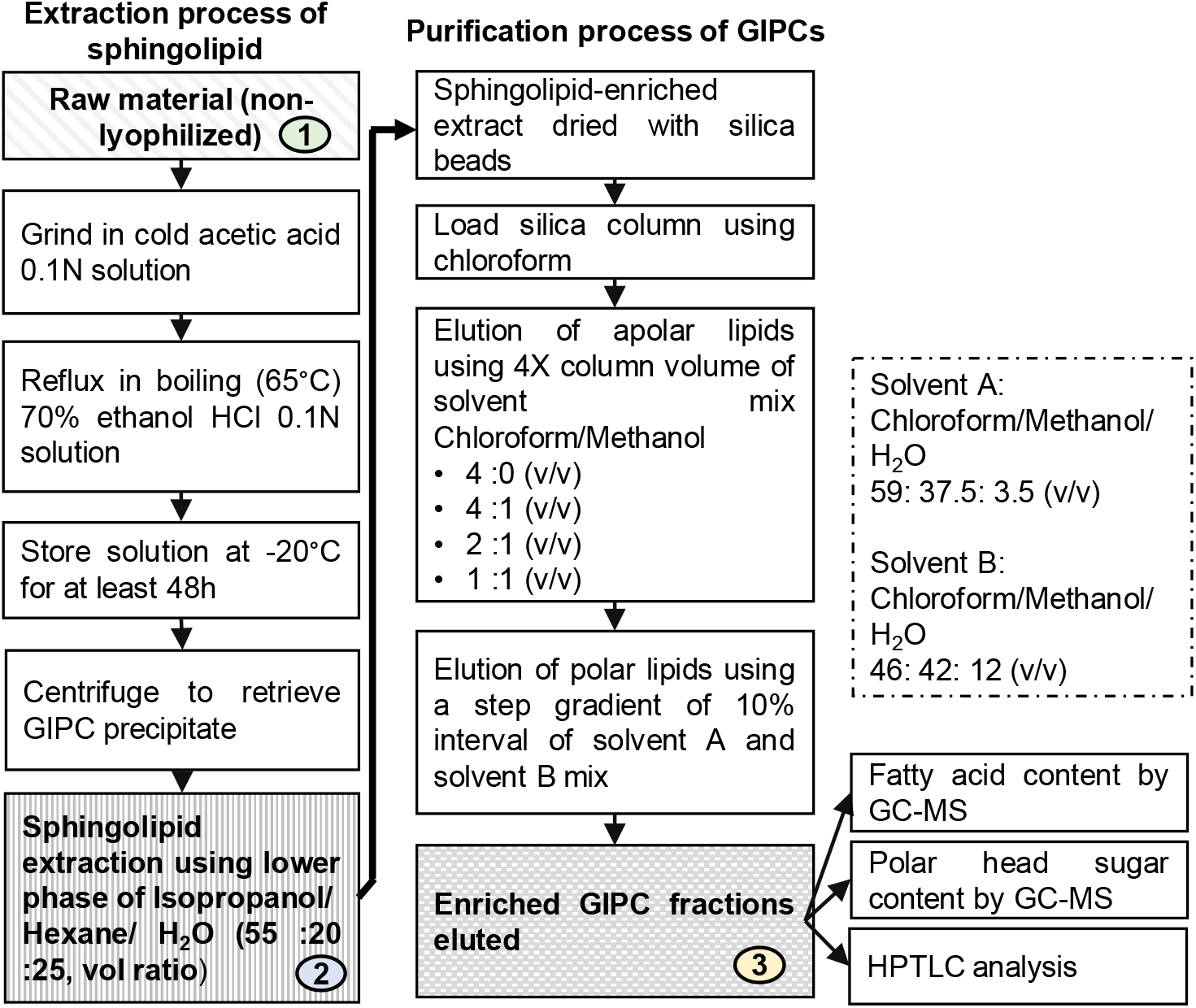
Extraction and purification protocol of GIPCs. GIPC purification scheme, adapted from (Carter & Koob, 1969; Kaul & Lester, 1975; Markham & Jaworski, 2007). The three steps labelled 1, 2 and 3, respectively are important milestones in the GIPC isolation steps.

### Fatty acid and sugar content of GIPC-enriched fractions

To estimate the GIPC content, as well as the phospholipid contamination (medium chain fatty acid FA of C16-18), samples were trans-esterified in hot methanol/sulfuric acid solution to release both fatty acid-esterified glycerolipids and fatty acid-amidified sphingolipids. The samples were then derivatized by trimethyl-silylation, and analysed by Gas Chromatograph-Mass Spectrometry (GC-MS) using internal standards, which allowed quantification of total FA content. The percentage of fatty acid in the samples with medium chain length (%FA) and hydroxylated and non-hydroxylated very long chain length (%(h)VLCFA) were calculated from 2 to 3 independent experiments. Samples retained after step 1 (raw plant material), step 2 (crude sphingolipid extract) and step 3 (GIPC-enriched fractions) were analysed for their fatty acid content (Figure 3A). As we proceeded through the purification steps, the amount of medium chain FA decreased as the amount of (h)VLCFA increased (Figure 3A). At step 2, the percentage of (h)VLVCFA in the sphingolipid extract was around 50% and at the final step, the amount of (h)VLCFA was at about 80% for all GIPC-enriched extracts (Figure 3A). The detailed FA composition of the GIPC-enriched fractions of all four species is provided in supplementary data 2. It was estimated that the enrichment in GIPC between the first and last steps of the extraction and purification process was 5-fold for Bo-GIPC, 4.2-fold for Nt-GIPC, 3.6-fold for Ap-GIPC but only 2-fold for Os-GIPC.

**Figure 3.**
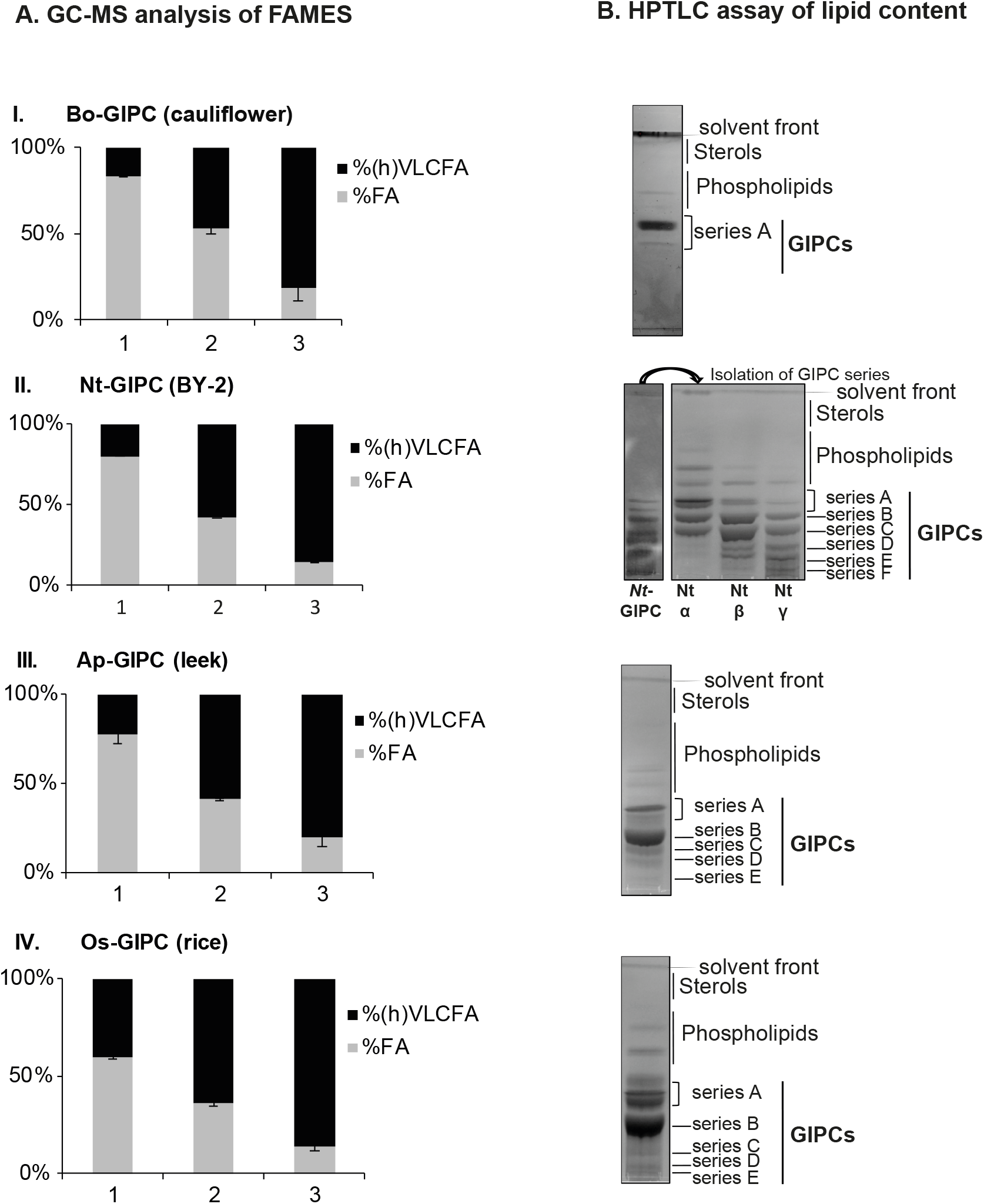
Gas Chromatography-Mass Spectrometry (GC-MS) analysis of fatty acid content after steps α, β and γ of the extraction and purification process (see Figure 2) (A) and High-Performance Thin Layer Chromatography (HPTLC) assay of lipid content after step 3 (B). A, Aliquots of (I) *Bo*-cauliflower, (II) *Nt*-BY-2, (III) *Ap*-leek and (IV) *Os*-rice samples at step 1,2 and 3 underwent trans-methylation to release fatty acid before derivatization by BSTFA, the resulting FAMES were analysed by GC-MS and the fatty acid content calculated (n=3). FA refer to fatty acid of 16 to 18 carbon atoms fatty acids and (h)VLCFA refer to hydroxylated or non-hydroxylated very long chain fatty acid of 20 to 28 carbon atoms. The amount of GIPC in each sample were extrapolated from the (h)VLCFA content. Data shown for 3 independent replicas. Error bars are SD. B, HPTLC assay shows the lipid content of GIPC-enriched samples after step 3. Cauliflower (I) contains mainly GIPC series A, BY-2 (Nt-GIPC) sample (II) were further separated by column chromatography to isolate the different GIPC series. Nt (α) GIPC-enriched sample contains mainly series A, B and C while Nt (β) and Nt(γ) show presence of polyglycosylated GIPCs (series D, E, F, etc). Leek (III) and rice (IV) samples contain mainly GIPC series B.

The final products were analysed by HPTLC to verify the lipid composition, and they revealed the predominance of GIPCs (Figure 3B). Only traces of sterols and phospholipids were observed, and glucosylceramide (GluCer) was not detected. As reported in (Buré et al., 2011), eudicots contained mainly series A, monocots series B and plant cells in liquid culture media a mix of GIPCs with highly glycosylated ones. The Bo-GIPC enriched fraction contained one major band of GIPC series A. The Nt-GIPC fraction contained GIPC series A to F, further separated into three fractions (α, β and γ) of increasing polarity. The less polar fraction α contained two bands of series A GIPC closely packed together, representing PhytoSphingoLipid 1, PSL1 (with *N*-acetyl glucosamine) and PSL2 (with glucosamine) as described in (Kaul & Lester, 1975) (Figure 3B), and a band of series B. The more polar fractions β and γ showed the presence of the highly polyglycosylated D to F series GIPC (Figure 3B). As previously published monocots, Ap-GIPC and Os-GIPC as enriched fractions contained mainly GIPC series B, and with some series A and polyglycosylated GIPCs also present (Figure 3B).

The predominant GIPC-derived (h)VLCFA species was dependent on the starting material. The Bo-GIPC enriched fraction consisted of h24, h24:1 and h26 as the main fatty acyl chain, Nt-GIPC with h24, h22, h23 and h25 acyl chain, Ap-GIPC with C24, h24, h22 and C22 and Os-GIPC with C24, C22, C20 and h24 (Figure 4A).

**Figure 4.**
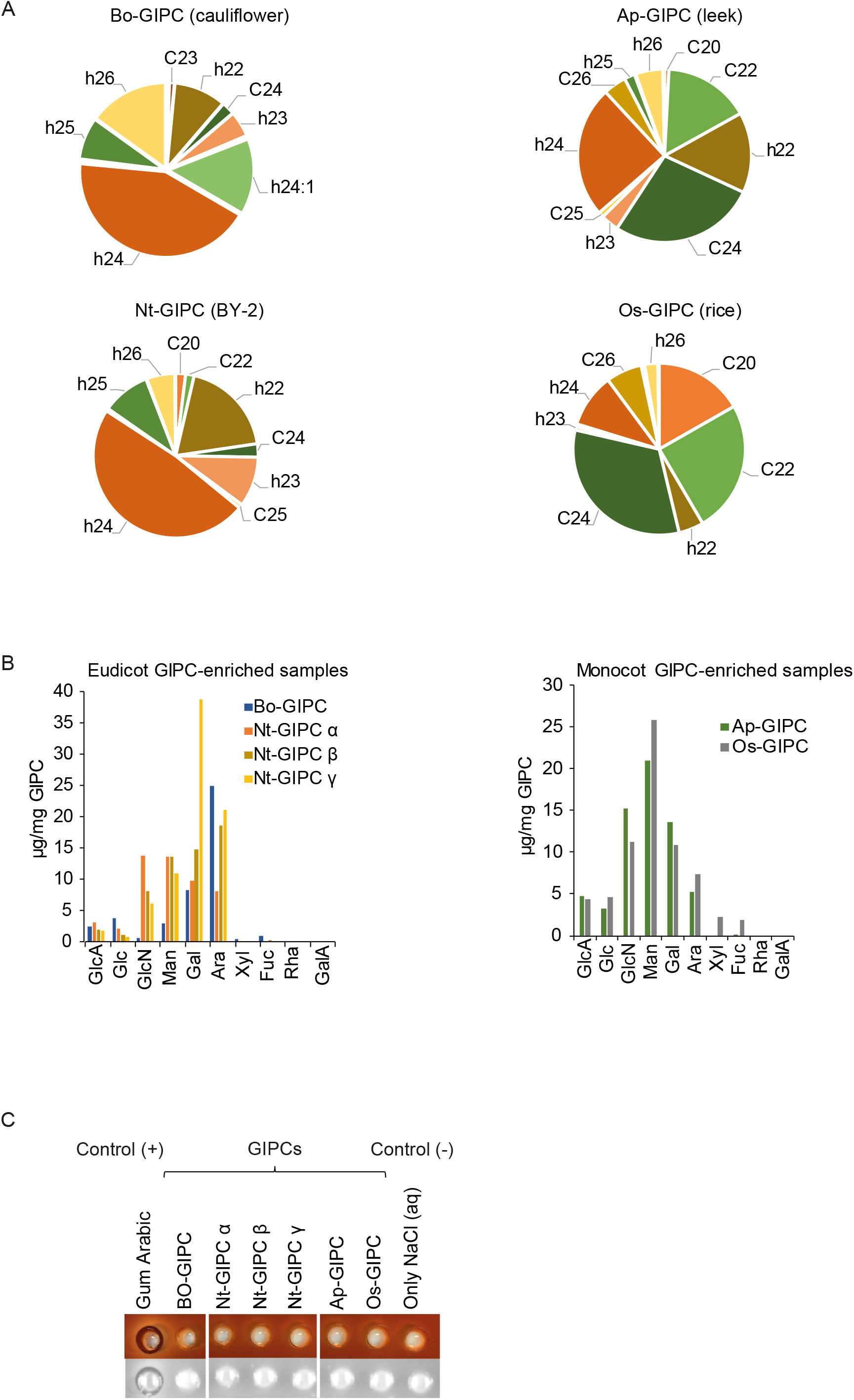
A, Very long-chain fatty acid (VLCFA) and hydroxylated VLCFA (hVLCFA) content of GIPC enriched samples from cauliflower, BY-2 cell culture, leek and rice cell culture. The fatty acids were released from the GIPC enriched samples by transmethylation followed by derivatization using BSTFA, before GC-MS analysis. B, HPAEC analysis of GIPC-enriched samples shows the monosaccharide content after TFA hydrolysis. Abbreviations are as follows: GlcA: glucuronic acid; Glc: glucose; GlcN: glucosamine; Man: mannose; Gal: galactose; Ara: arabinose; Xyl: xylose; Fuc: fucose; Rha: rhamnose; GalA: galacturonic acid. C, Yariv reactivity test of GIPC-enriched samples to detect arabino-galactan content. No arabino-galactan were detected. 50ug of each sample (1mg/ml) was deposited in each well, the picture was taken 48h after initiating the reaction.

We next investigated the sugar moieties present in GIPC-enriched fractions by high performance anion exchange chromatography (HPAEC) coupled with Pulsed Amperometric Detection (PAD), a technique used to detect underivatized monosaccharide sugars (Figure 4B). Since different glycosidic bonds hydrolyse at different rates, and some sugars rings are more sensitive to cleavage than others, the sugar content of the Bo-GIPC enriched fraction was analysed 1h, 3h and 4h after trifluoroacetic acid (TFA) hydrolysis. Results showed that hydrolysis has no to very little effect on the sugar moieties of GIPC fractions (Supplementary data 3). As expected, all GIPC-enriched fractions contained glucuronic acid (GlcA) found in GIPC samples previously characterized, see for review (Mamode Cassim et al., 2019). The Bo-GIPC enriched fraction not only contains glucose (Glc) and mannose (Man) previously found in Brassicaceae species (Fang et al., 2016), but also large amount of arabinose (Ara) and galactose (Gal). These latter sugars, never described in Brassicaceae such as Arabidopsis, could be a real specificity of Bo-GIPC’s polar head, or due to cell wall/glycoprotein contamination of the GIPC-enriched fraction, see further below.

The different fractions of the Nt-GIPC series have a complex glycan content. Fraction α contained glucuronic acid (GlcA), glucosamine (GlcN) and mannose (Man) (Figure 4B). Note here that *N*-acetyl glucosamine (GlcNAc) is hydrolysed during the extraction procedure and is mixed with GlcN. Gal and Ara became the main glycan moieties in fractions β and γ as described for highly glycosylated GIPC, series D (Kaul & Lester, 1978). Monocot GIPC-enriched sample, both Ap-GIPC and Os-GIPC contained mainly Man, Gal and GlcN at relatively equal amount, and GlcA and Ara at lower amount (Figure 4B).

Previous studies have suggested interactions between GIPCs and cell wall components, particularly the pectin Rhamnogalacturonan II (RGII) (Voxeur & Fry, 2014). However, we did not detect either galacturonic acid (GalA) nor rhamnose (Rha), two main components of pectins, suggesting no major pectin contamination (Figure 4B). We detected, however, a large amount of Ara and Gal (Figure 4B). A Yariv reactivity test (Kitazawa et al., 2013) was performed to check for the presence the arabino-galactan (AG) as contaminants in the GIPC-enriched fractions (Figure 4C). No zone of clearance was observed, suggesting no detectable AG in each GIPC sample (50 μg). Gum arabic and saline buffer were used as positive and negative controls respectively (Figure 4C). The potential contamination of the GIPC samples by proteins was also tested using the Bradford method. However, in GIPC samples of up to 30 μg, no protein was detected (data not shown).

The Bo-GIPC and Ap-GIPC purified fractions were analysed by LC-MS and compared to total sphingolipids extracted from crude cauliflower or crude leek. Results showed in Supplementary data 4 revealed an absence of CER and gluCER contamination in the purified GIPC sample, and that the LCB and FA content is very similar except a slight loss of h24:0/1- and t18: 0/1-containing GIPC (less that 10%), see Supplementary data 5.

### Biophysical characterisation of the GIPC-sterol interaction

We decided to focus on the Bo-GIPC to perform various biophysical analyses. We first characterized the lipid-lipid interactions at the micrometric level by the Langmuir trough compression technique applied on a monolayer model at the air-water interface (Deleu et al., 2014). As previously reported (Cacas et al., 2016), experimental biophysical characterization coupled to energetic calculations suggested a preferential interaction of GIPC series A with β-sitosterol, defined as a ‘condensing effect’, with the favourable interaction minimizing the energy of interaction, i.e. where the interfacial area occupied by the 2 molecules is less than the interfacial area occupied by single one. To further characterize the interaction of Bo-GIPC with phytosterols, we conducted biophysical experiments to investigate the outer leaflet organization with free and conjugated sterols (β-sitosteryl Glucoside, SG and Acyl (18:2) β-sitosteryl Glucoside, ASG). The ratio of GIPC:sterol (80:20 mol ratio) is consistent with the estimated ratio of the lipids in the outer leaflet of the PM (Cacas et al., 2016)(Tjellstrom et al., 2010). The compression isotherm of Bo-GIPC (green line) (Figure 5A) shows a low and relatively constant surface pressure in large molecular areas, corresponding to a ‘gaseous’ state. Compression of the monolayer induced a progressive increase in surface pressure, indicating the appearance of a liquid-expanded state (in agreement with the two-dimensional compressibility modulus, Cs^−1^, of 38.3 mN m^−1^ in the 160-to 110-Å^2^ per molecular region), which is characterized by a certain degree of condensing interaction between the molecules at the interface (Figure 5A). The mean interfacial area of Bo-GIPC is 212.9 ± 4.9 Å^2^ in its expanded form and at its most condensed form is 60.0 ± 14.6 Å^2^. These results are in agreement with the results previously obtained with *Nicotiana tabaccum*-GIPCs (Cacas et al., 2016).

**Figure 5.**
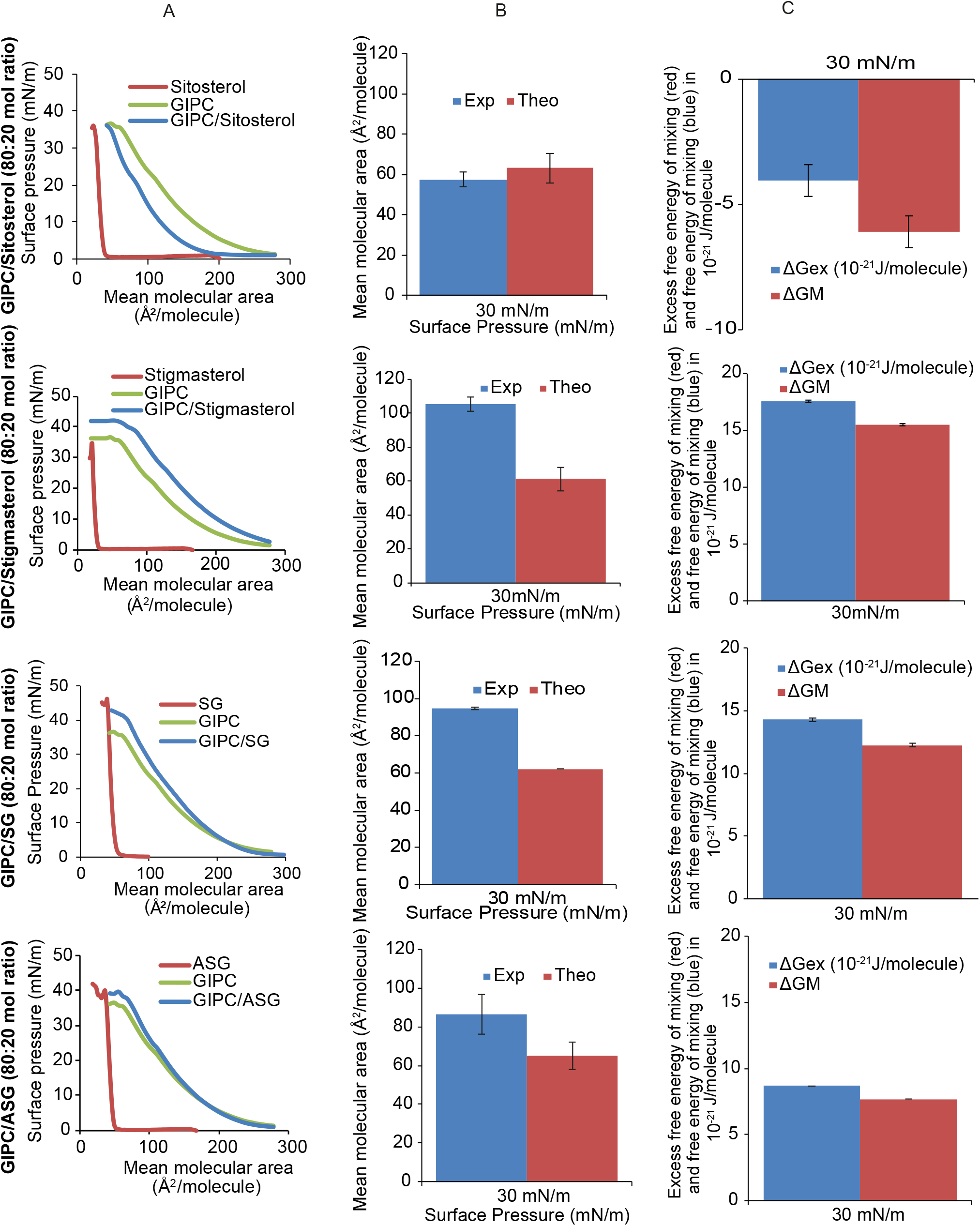
Surface pressure-area (π-A) isotherms, at the air-aqueous phase interface, of pure GIPC and sterol monolayers and of mixed GIPC/sterol monolayer prepared at a molar ratio of 0.80. A. The isotherms were recorded at 25°C on an aqueous subphase composed by 10 mM Tris buffer at pH 7. Duplicate experiments using independent preparations yielded similar results. B, Comparison of the experimental (blue bars) and theoretical (red bars) mean molecular areas at a surface pressure of 30 mN/m for a GIPC/sterol molar ratio of 0.80. The theoretical value is obtained according to the additivity rule: A_12_ = A_1_X_1_ + A_2_X_2_, where A_12_ is the mean molecular area for ideal mixing of the two components at a given π, A_1_ and A_2_ are the molecular areas of the respective components in their pure monolayers at the same π, and X_1_ and X_2_ are the molar ratios of components 1 and 2 in the mixed monolayers. C, Excess free energy of mixing (ΔGex; blue bars) and free energy of mixing (ΔGM; red bars) of the mixed monolayer GIPC/sterol at a molar ratio of 0.80 at the surface pressure of 30mN/m. ΔGex and ΔGM were calculated according to the equations as shown in (Eeman et al., 2005; Maget-Dana, 1999). Abbreviations are as follows: SG, steryl glucoside (sitosterol, glucose head group); ASG, acyl steryl glucoside (sitosterol, glucose head group and C18:2 acyl chain). Error bars are SD.

The interaction of Bo-GIPC mixed with different sterols was assessed by the thermodynamic analysis of the compression isotherms of mixed GIPC-sterol monolayers. In this comparative study, we adhere to the rule of additivity, which suggests that if two molecules within a mixed monolayer are immiscible, the area occupied by the mixed film will be the sum of the areas of the separated components. The deviation to that rule is attributed to the existence of specific interaction between the two molecules (Maget-Dana, 1999). The mean molecular area of the mixed monolayer Bo-GIPC: β-sitosterol (80:20) was lower than the calculated theoretical value (using the rule of additivity), at the estimated physiological membrane surface pressure of 30 mN m^−1^ (Marsh, 1996) (Figure 5B). This condensing effect of β-sitosterol in presence of Bo-GIPC confirms that previously reported for tobacco GIPCs (Cacas et al., 2016). The trend was however reversed for the mixed monolayers of Bo-GIPC:SG (80:20) and Bo-GIPC:stigmasterol (80:20), where the mean molecular area is significantly higher than the theoretical value (Figure 5B). For Bo-GIPC:ASG (80:20), the effect is intermediate. The most significant difference between the experimental and theoretical mean molecular area was obtained for the mixed monolayer Bo-GIPC:stigmasterol (80:20). Interestingly, the only structural difference between β-sitosterol and stigmasterol is the presence of a double bond at C22 in stigmasterol. The mixed monolayer GIPC:ASG (80:20) had a comparable mean molecular area to GIPC molecule at low surface area (Figure 5A) and the average difference between the mean molecular area and its theoretical value is 30 Å^2^ per molecule for all three surface pressures (Figure 5B).

In order to thermodynamically analyse the interaction of the two components and the stability of the mixed monolayer, the excess free energy of the mixing (ΔGex) and the free energy of mixing (ΔGM) were respectively calculated for all four mixed monolayers (Figure 5C). The negative value of ΔGex for the mixed monolayer GIPC: β-sitosterol (80:20) suggested a strong attractive interaction between the two components and the negative value of ΔGM indicated thermodynamic stability of the mixed monolayer (Figure 5C) as suggested by (Cacas et al., 2016). The values of ΔGex and of ΔGM for the mixed monolayers GIPC:SG (80:20), GIPC:ASG (80:20) and GIPC/Stigmasterol (80:20) were both positive in all three mixed monolayers (Figure 5C) showing repulsion between the molecules within the monolayer and thermodynamic instability of the mixed monolayers.

### Modelling the interaction between GIPC and phytosterols

Hypermatrix is a simple docking method used to calculate specific interactions between two amphiphilic molecules (for a review see (Deleu et al., 2014). The interaction of one molecule of GIPC series A with t18:0/h24:0 and one molecule of sterol was generated *in silico* using this method and analysed. The sterols used were the four molecules studied by the Langmuir monolayer technique i.e. β-sitosterol, stigmasterol, ASG, and SG (Figure 6). The interacting molecules displayed very different configurations. The differences between the spatial organization of the GIPC/sitosterol and GIPC/stigmasterol were striking: the α-side of the steryl moities of β-sitosterol was directed towards the acyl chains of the GIPC, whereas the steryl rings of stigmasterol was positioned at a perpendicular angle with respect to GIPC hydrocarbon chains (Figure 6A,B). In mammals the interaction of the α face of cholesterol with lipid acyl chains favours its condensing effect (Rog & Pasenkiewicz-gierula, 2004). This was notably established by comparing the effects of cholesterol and lanosterol, which possesses a methyl group on the α face, on lipid organization (Yeagle et al., 1977)(Smondyrev & Berkowitz, 2001)(Róg et al., 2009). For stigmasterol, the structural difference of the unsaturation on C22 seems thus to modify its interaction with GIPC (Figure 6B), and this can be correlated to the non-condensing effect observed experimentally in the monolayer compression experiments (Figure 5). Similarly, the β face of the steryl ring moiety of the steryl glucoside (SG) was oriented towards GIPC acyl chains. It is noteworthy that the bending of the sugar head group of GIPC favours its interaction with the glucose head group of SG (Figure 6C). In the conformation of the conjugated sterol ASG, the acyl chain is in direct interaction with the α side of the sterol, such that the β-side of the steryl cycle interacts with GIPC acyl chains (Figure 6B). Thus, β-sitosterol is the only sterols tested for which the interaction of its α-face with the GIPC acyl chains is favoured. This could be related to its condensing effects observed experimentally.

**Figure 6.**
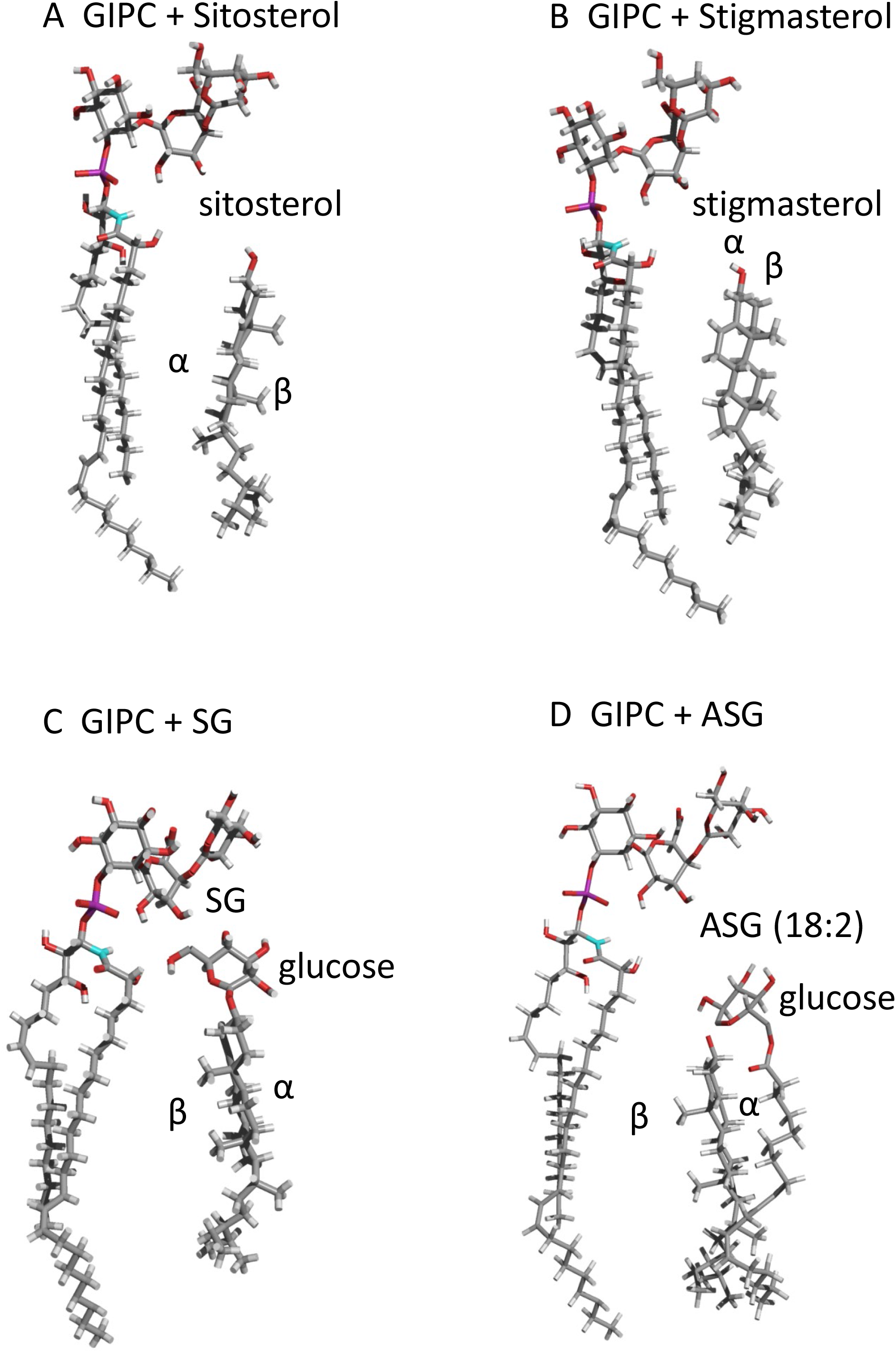
Modelling of the interaction of GIPC and sterols. Theoretical interactions calculated by HyperMatrix docking method with one molecule of GIPC serie A t18:0/h24:0 and one molecule of either A, β-sitosterol, or B, Stigmasterol or C, Steryl Glucoside, SG (β-sitosterol, with glucose head group), or D Acyl Steryl Glucoside, ASG (β-sitosterol, with glucose head group/18:2 acyl chain).

### Effect of GIPC on membrane organization and thickness

To further investigate the properties of GIPC using model bilayers, we tried to make large unilamellar vesicles (LUV) with sizes between 50-500 nm with Bo-GIPC by freeze/thawing. We found that GIPC alone made aggregates but not vesicles (Figure 7A). Fluorescent microscopy observations of Nt-GIPC containing LUV in water at RT, pH 7 led to similar results (Supplementary data 6A). However, by adding phospholipids we could observe the formation of LUV (Supplementary data 5A). To closer mimic the outer leaflet of the PM enriched in GIPCs, we generated LUV with a ternary system of GIPC:phospholipid: β-sitosterol (1:1:1). As phospholipids, we used 1-palmitoyl-2-linoleoyl-*sn*-glycero-3-phosphocholine (PLPC), 1-palmitoyl-2-oleoyl-*sn*-glycero-3-phosphocholine (POPC) or dioleoyl-*sn*-glycero-3-phosphocholine (DOPC), in all cases the ternary system yielded liposomes using the freeze-thaw method with liquid nitrogen and water bath of 60°C (Figure 7A) (Supplementary data 6B). Giant unilamellar vesicles (GUVs) were also made using the Teflon method by (Kubsch et al., 2017) with a ternary mix of GIPC/DOPC/sitosterol (Figure 7B). The incorporation of GIPCs into the liposomes were analysed by dynamic light scattering (DLS) which gives the hydrodynamic diameter of the liposomes. The addition of GIPC did not seem to affect the hydrodynamic diameter of liposomes, which was about 100 nm. The ζ-potential of the GIPC-containing liposomes was measured to be around −26 mV, while DOPC/ β-sitosterol alone had a ζ-potential of −5 mV (Figure 7C). The difference in ζ-potential between GIPC and GIPC-free liposomes is attributed to the fact that GIPCs are negatively charged because of the presence of the glucuronic acid, and furthermore, confirms that GIPC was indeed incorporated into the lipid membrane. (Jiang et al., 2019) showed that the ζ-potential of the surface of wild-type *Arabidopsis* mesophyll cells are at −20mV which is quite close to that of our liposomes. It seems that GIPC contributes significantly in the negative potential of the plant PM outer leaflet, which might influence, for example, its interaction with cell wall components. The structure of the LUVs made by the freeze/thaw method was further investigated by cryo-EM. GIPC/POPC-^2^H_31_/ β-sitosterol- and GIPC/POPC-^2^H_31_/stigmasterol-containing LUVs formed not only regular-shaped bilayer vesicles, but also planar bilayer structures that seem more rigid and not able to bend and make proper vesicles (arrow in Figure 8). Comparison of the bilayer thickness of these GIPC-containing LUVs with LUVs containing only POPC and sterols, showed a significant difference of thickness from 4.5 nm for the ternary LUV to 3.5 for the binary LUV (Figure 8).

**Figure 7.**
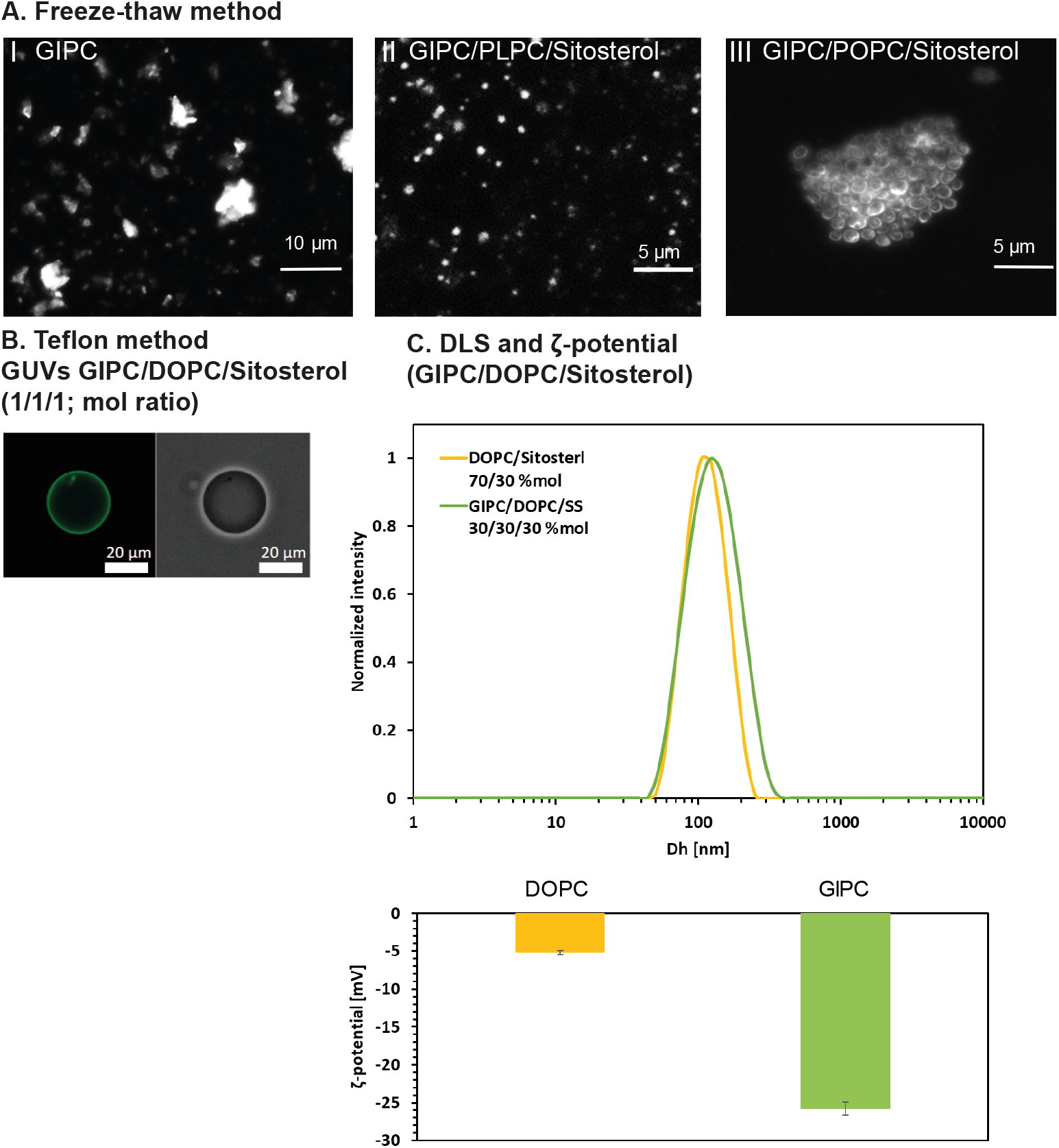
A, Bo-GIPC containing-liposomes in buffer solution after 3 cycles of freeze and thaw. Enriched Bo-GIPC (cauliflower), underwent freeze (−20°C, 20 min) and thaw (60°C, 20 min) cycles three times GIPC in TBS buffer pH 5,8 with or without phospholipid and β-sitosterol at a concentration of 1 mg/ml. (I) GIPCs alone form crystals in a saline buffer solution. A lipid mix, at a concentration of 1mg/ml, of GIPC/PLPC/ β-sitosterol or GIPC/POPC/ β-sitosterol (1:1:1, mol/mol), shown in (II) and (III) respectively, forms vesicles of approx. 2 μm. B, Fluorescence and phase contrast microscopy images of Giant unilamellar vesicles (GUVs) of GIPC/DOPC/ β-sitosterol (1:1:1, mol/mol). The lipid mix was labled by NBD-PC at 0.1%. mol. C, Dynamic light scattering (DLS) and ζ-potential of liposomes containing DOPC/ β-sitosterol (7:3, mol ratio) (yellow) and GIPC/DOPC/ β-sitosterol (1:1:1, mol ratio) (green), respectively provide the size which is around 100nm and ζ-potential values of −28 mV in the presence of GIPC. Error bars are SD.

**Figure 8.**
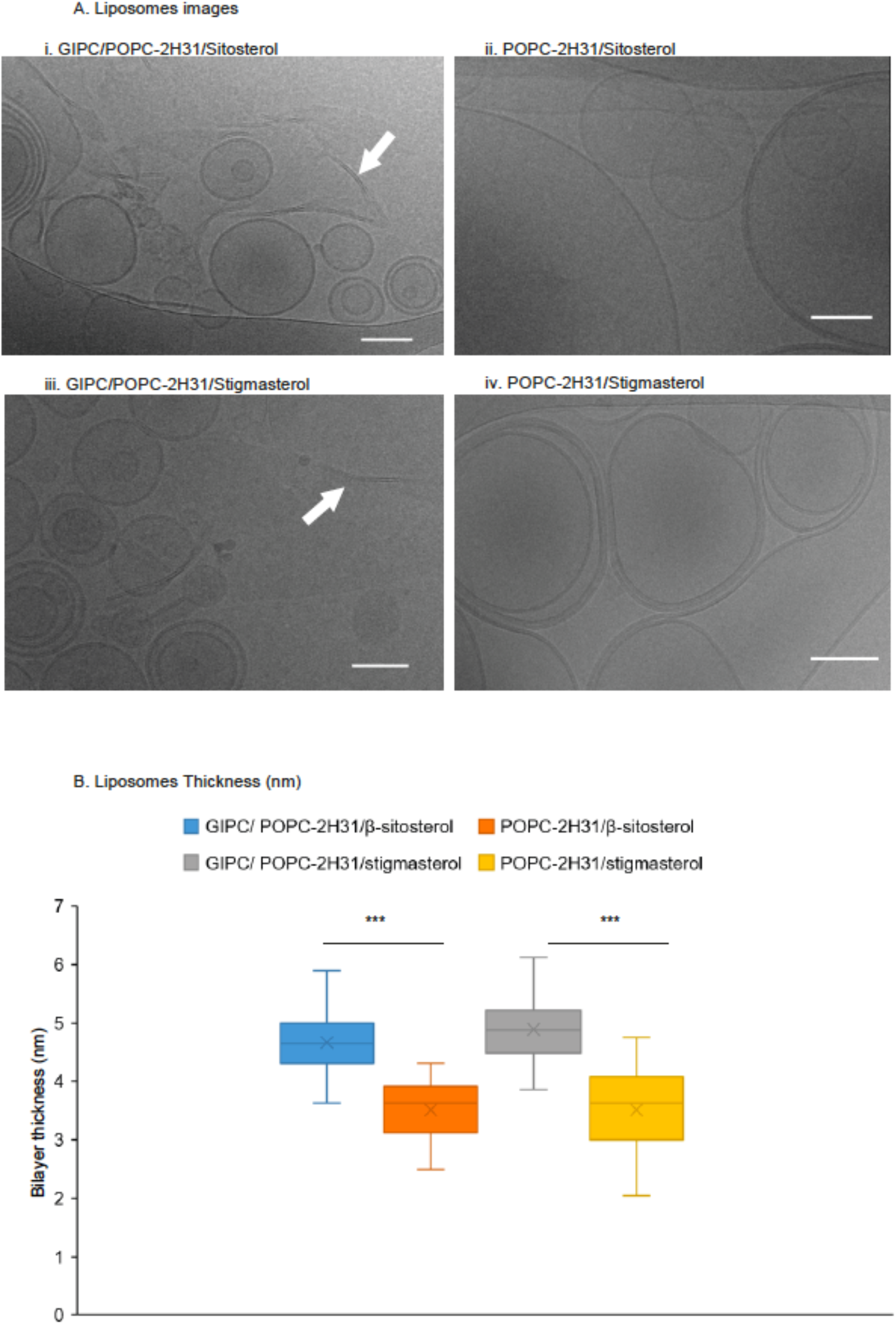
Liposomes shapes varies with lipid composition. A, Cryo-EM images of liposomes. POPC-d31 that is a deuterated POPC on the carbon of the palmitoyl chain: 16:0-d31-18:1 PC, in the presence of sterols (ii sitosterol and iv: stigmasterol) are mainly present as vesicles, showing one to few bilayers. In the presence of GIPC, these liposomes are still observed but at the same time, rigid bilayers structures appearing as flat entities are also observed (white arrows in I and iii). Scale bar, 100 nm. B. membrane thickness measurements. Measurements were made using ImageJ software to compare the membrane thickness with or without GIPC. Error bars are SD. Significance was determined by Student’s *t*-test. ***P < 0.0001

We also investigated the influence of GIPCs on membrane thickness by neutron reflectivity (Figure 9A). Supported Lipid Bilayers (SLB) were formed by vesicle fusion of liposomes containing POPC, GIPC and constant β-sitosterol concentration. Three different membrane compositions were tested with increasing (0, 15 and 30% mol) GIPC concentration. GIPC. The reflectivity profile was analysed, and following model fitting, the scattering length density profile and the thickness of the polar head and acyl tail in the bilayer were obtained. The results showed that liposomes containing 30% mol. of GIPC did not form a continuous bilayer on the surface, as indicated by a high solvent content in the hydrophobic tail region. This implies that the high GIPC content modified the bilayer properties, such that it did not adhere to the support. However, 0 and 15% mol containing GIPC liposomes did form continuous bilayers. The addition of GIPCs increased the bilayer thickness by 8 Å, as compared to GIPC-free SLBs, due to the 4 Å of sugar head group in each layer (Figure 9A). Refer to the tables of Figure 9B for more details of the structural parameters that were generated. Figure 9C shows a scattering length density profile of the SLBs.

**Figure 9.**
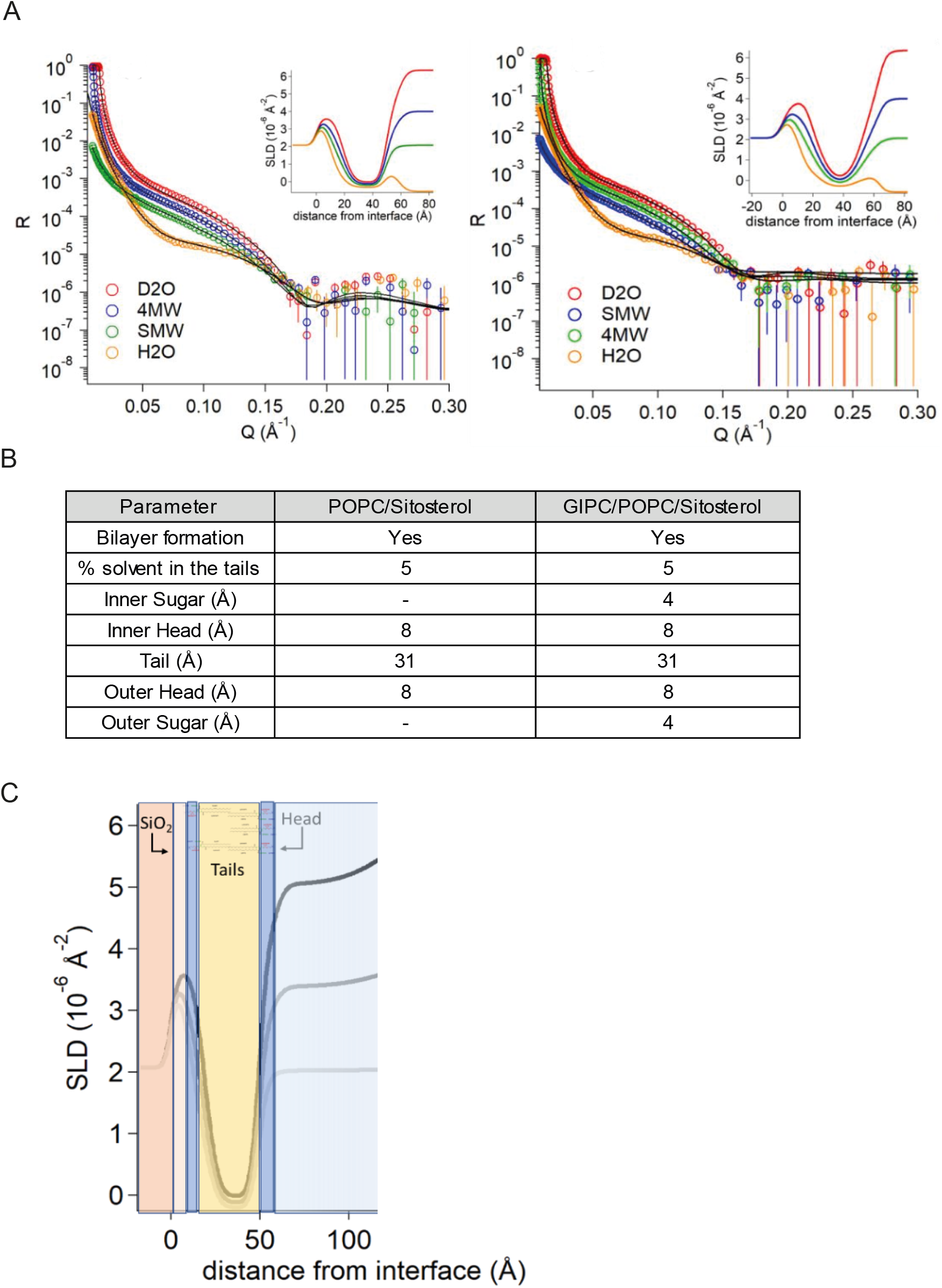
Reflectivity profiles and calculated Scattering length density (SLD) following lipid bilayer deposition of (i) POPC/ β-sitosterol (70:30, mol/mol) and (ii) GIPC/POPC/ β-sitosterol (55:15:30, mol/mol). A, The multilayer model was composed from the silicon substrate (SLD=2.07 10^−6^Å^−2^) covered with a layer of silicon oxide (SLD=3.47 10^−6^Å^−2^). B, Structural parameters after multilayer model fitting of reflectivity profiles of lipid bilayer. C, Scheme showing the SLD profile overlaid on the multilayer model as obtained for POPC membrane.

### Phase transition of GIPC containing-vesicles analysed by liquid state ^2^H-NMR

Finally, using solid-state ^2^H-NMR, we asked whether GIPCs have an effect on the gel-to-fluid phase transition of a fully hydrated binary and ternary lipid mix system. Solid-state ^2^H-NMR spectroscopy is a non-intrusive method giving structural and dynamic information about lipid bilayers (Davis, 1983; Seelig, 1977). Here, we aimed to find the nature of the membrane phases, their dynamics, and how GIPCs and phytosterols are regulating the membrane phase transition, such as the well-described effect of cholesterol on the membrane (Oldfield et al., 1978).

We used Bo-GIPC enriched fractions to make membrane systems using deuterated palmitoyl-oleoyl phosphatidylcholine containing 31 atoms of deuterium on the palmitoyl chain (POPC-^2^H_31_) as a probe for solid-state ^2^H-NMR. We chose to use POPC as it is a phospholipid with a long chain fatty acid with an unsaturation found in plant PM (Cacas et al., 2016) and a gel-to-fluid transition temperature of −2.5°C ± 2.4 (Koynova & Caffrey, 1998). We generated liposomes using the freeze/thaw method as previously described. Figure 10A showed ^2^H NMR spectra of two lipid mix systems containing GIPC (GIPC/ POPC-^2^H_31_/β-sitosterol (1:1:1, mol ratio) and GIPC/ POPC-^2^H_31_/stigmasterol (1:1:1, mol ratio) and two control samples without GIPC (POPC-^2^H_31_/β-sitosterol (1:1, mol ratio) and POPC-^2^H_31_/stigmasterol (1:1, mol ratio). Spectra were acquired by varying the temperature from −10°C to 40°C. These are plausible thermal variations that plants may experience in nature. The obtained ^2^H NMR spectra exhibit the typical powder pattern line shape with a spectral width decreasing as the temperatures increase. This qualitative observation can be supplemented by a quantitative characterization using the first spectral moment (Davis, 1983). Figure 10B shows the temperature plots of first moments (M1) calculated from ^2^H-NMR powder spectra of liposomes with or without Bo-GIPCs, as well as pure POPC-^2^H_31_. On figure 10B left, we can hence appreciate the phase transition of a pure POPC-^2^H_31_ membrane such that the low M1 corresponds to the fluid (*Ld*) phase and the high M1 to the rigid (*Lo*) phase. The thermal variation showed an abolished phase transition upon adding phytosterols to POPC-^2^H_31_ (Figure 10B left). This abolition is more pronounced for β-sitosterol with a higher ordering effect above the phase transition temperature compared to stigmasterol. These conclusions can be transposed to ternary systems with the difference that β-sitosterol has a stiffening effect at low temperatures (Figure 10B right). Above the POPC-^2^H_31_ phase transition both GIPC and phytosterol were able to rigidify the membrane, with a larger effect for β-sitosterol. This result is similar to those obtained by Beck et al, 2007. Taken together, these experiments showed that GIPC and phytosterols adopt the same behaviour as cholesterol, and hence have a high propensity to regulate fluidity during temperature variations.

**Figure 10.**
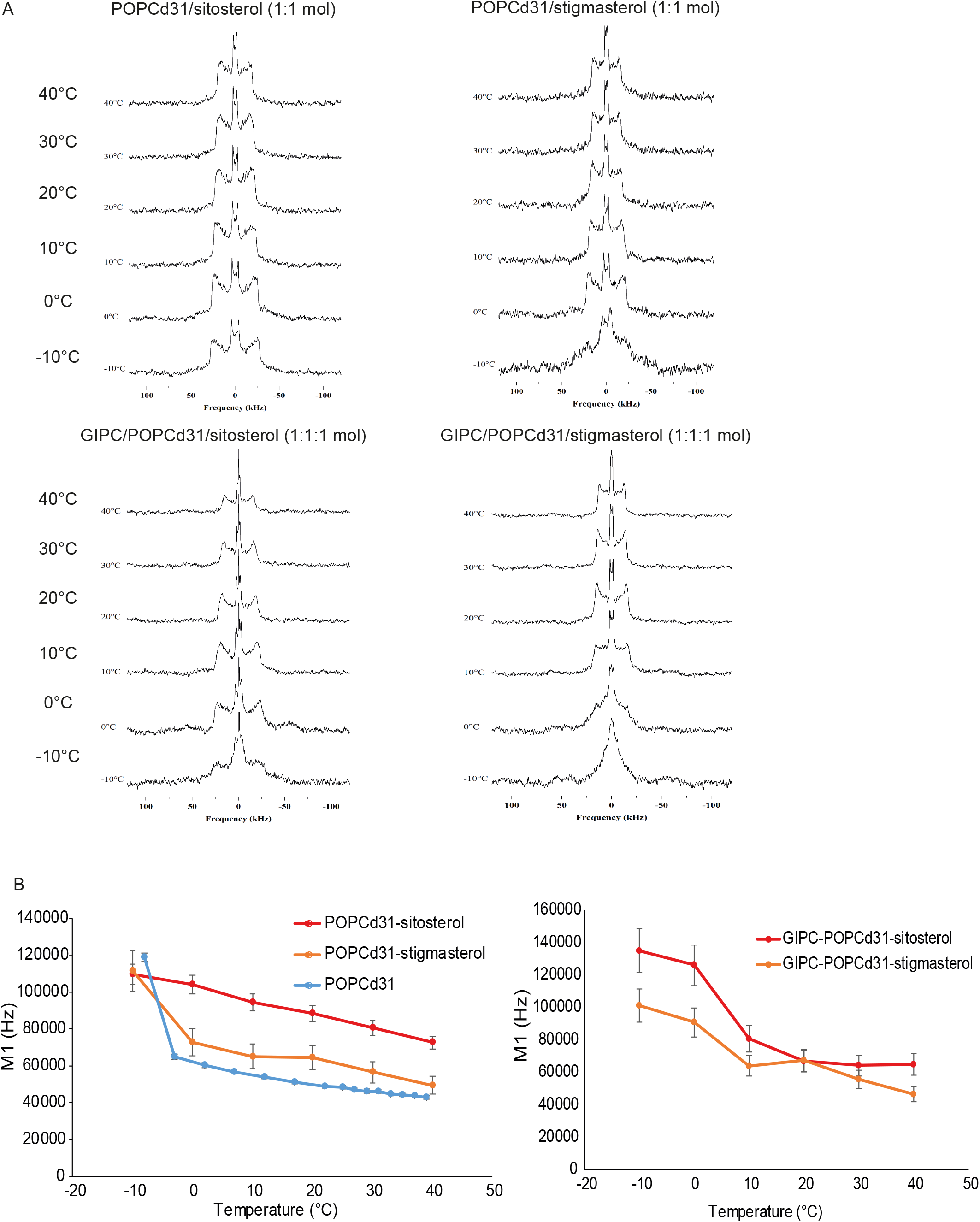
A, ^2^H-NMR powder spectra of lipid mix and B, First spectral moment of ^2^H-NMR spectra showing membrane ordering *vs.* temperature POPC-^2^H_31_system alone, in binary systems of POPC-^2^H_31_/β-sitosterol (1:1 mol/mol) and POPC-^2^H_31_/Stigmasterol (1:1 mol/mol) and ternary systems of GIPC/ POPC-^2^H_31_/ β-sitosterol (1:1:1 mol ratio) and GIPC/ POPC-^2^H_31_/Stigmasterol (1:1:1 mol ratio).

## Discussion

### Fine-tuning GIPC purification

GIPCs are quantitatively and qualitatively essential components of plant plasma membranes (Cacas et al., 2013; Mortimer et al., 2013). As there are no commercially available GIPCs, reasonable quantities of these molecules must be isolated with good purity to study their biophysical properties. This paper describes a protocol for efficient purification of milligram amount of GIPCs. It has been inspired by three publications (Carter & Koob, 1969),(Kaul & Lester, 1975)(Markham & Jaworski, 2007), whereby the steps were rearranged in order to get rid of contamination such as sterols and glycerolipids during the extraction procedure, while retaining GIPCs which were eluted in the column purification steps. This is simpler and more convenient than the one used in previous publications where large amounts of solvents were used. It is also time-efficient and achieve a reasonable yield since we were able to obtain 30 mg of GIPC of up to 85% purity from 4 distinct plant materials of 600-800g fresh weight. In the near future, to improve yield and purity, we still have to fine-tune the purification and extraction process of GIPCs, and it will also be important to purify different GIPC series individually, so as to decipher the number, bonds and types of sugar residues which make up the different plant GIPC polar head groups, as was done for fungi GIPC by NMR (Simenel et al., 2008) (Gutierrez et al., 2007). To do so, *ad hoc* preparative chromatography needs to be developed with elution solvents with the right polarity.

The full structure and diversity of sugar moieties in the GIPCs polar head remains to be understood and investigated. The diversity seems to be important, for example in plant/pathogen interactions. A recent study showed that the GIPC polar head may be receptor for oomycete necrotic toxins called Necrosis and ethylene-inducing peptide 1– like (NLPs). Plants enriched in GIPC series A are sensitive to NLPs while those enriched in GIPC series B are insensitive to NLPs, hence conferring resistance against pathogens secreting NLPs (Lenarčič et al., 2017).

The polar head sugar diversity of GIPCs are clearly species-dependent (Buré et al., 2011). For instance, the increasingly large amounts of Gal and Ara in the Nt-GIPC fractions has been described in (Kaul & Lester, 1978) where GIPCs of tobacco leaves contain up to 4 Ara and 2 Gals attached to the core GIPC structures of GlcN/GlcNAc-GlcA-Ins-P-Cer. Hence, it is correct to assume that the large amount of Ara and Gal of Nt-GIPC(fraction γ) derives from the polyglycosylated GIPCs of up to GIPC series E of Ara-Ara-Gal-Gal-Man-GlcN/GlcNAc-GlcA-Ins-P-Cer.

In all enriched GIPC fractions purified in this study, GalA and Rha are absent (Figure 4B), suggesting no contamination by pectins, particularly RGII, which is reported to bind to GIPCs (Voxeur & Fry, 2014). It should be noted that Ara and Gal can form AG side chains of the pectin Rhamnogalacturonan I (RGI). Ara and Gal are also the dominant sugars in the abundant GPI-anchored plasma membrane glycoproteins, the arabinogalactan proteins (AGPs). To test for the presence of RGI or AGPs, we performed a Yariv reactivity test, but we did not detect anything that could bind to Yariv.

Previous work in *Arabidopsis thaliana* (*At*, also a Brassicaceae, like cauliflower), identified a mannose as the first sugar attached to the GlcA-Ins-P-Cer core in vegetative tissues, a reaction catalysed by the glycosyltransferase *AtGMT1* (Fang et al., 2016). We hypothesized that this also might be the case in cauliflower. In the seed tissue of *At*, Glucosamine InositolPhosphorylCeramide Transferase 1 (*AtGINT1*), another glycosyltransferase, adds a GlcNAc instead of a Man to the core structure of GIPC (Ishikawa et al., 2018). However, in both cases, the authors also detected Ara and Gal as part of the sugar composition of their GIPC enrichments. Therefore, we cannot yet conclude whether Ara and Gal are contaminations or are inherent sugar moieties of GIPC.

### GIPCs alone form aggregates, but not liposomes

With the aim of studying the physical properties of GIPC, we attempted to make liposomes (LUV). Since GIPCs have large polar heads and VLCFAs, they tend to agglomerate, similarly to pure sterols. In order to make liposomes and further study GIPCs, a binary mix with phospholipids, or a ternary mix with phospholipids and sterols were used. To be closer to the biological PM model, we used a lipid mix of GIPC/phospholipid/sterol at a molar ratio of 1:1:1. As expected for lipids with (h)VLCFAs, GIPCs increase the thickness of the model membrane by a few nm, as shown by neutron reflectivity on a supported bilayer. The bilayer thickness of liposomes containing GIPCs as observed using cryo-EM are around 6-7 nm for the ternary mix, which corresponds well with the observed thickness of purified PM from *Medicago truncatula* and tobacco (Lefebvre et al., 2007)(Mongrand et al., 2004).

One important feature of the PM is its electrostatic charge. PM purification using polymers phase separation PEG/Dextran relies on the fact that PMs are highly negatively charged and that the PM right-side-out (RSO) fraction is attracted to the positively charged PEG phase (Morré & Morre, 2000). The membrane surface charge (MSC) is regulated by lipids and post transcriptional modification of proteins such as phosphorylation (Goldenberg & Steinberg, 2010). The ζ-potential of GIPC-containing liposomes is −26 mV, five times higher than a PC/ β-sitosterol-containing bilayer, likely due to the large negativity of GIPC conferred by its phosphate group and the GlcA residue of the polar head. Therefore, since GIPCs are mostly located in the outer leaflet of the PM (Cacas et al., 2016), we conclude here that GIPCs contribute strongly to the negative charge of the PM outer leaflet.

### GIPC interaction with sterols and effects on the membrane order

Using the biophysical techniques of Langmuir compression isotherms and molecular modelling, we showed that GIPCs interact differentially with different phytosterols. We confirmed that GIPCs with β-sitosterol has a condensation effect as described in (Cacas et al., 2016) whereas non condensing interactions occur between stigmasterol, SG or ASG and GIPCs (Figures 5, 10). These differential interactions appear to be structure dependent. Just adding a glucose head group (SG) and an acyl chain (ASG) or an unsaturation (C22 in stigmasterol) to the β-sitosterol steryl moieties changes the interaction with the GIPCs and modifies the properties of the model membranes (Figure 5, 6 and 7). Interestingly, GIPCs, ASG and SG all accumulate after drought stress (Tarazona et al., 2015). The differential interaction of GIPCs with the different kind of sterols could explain how plants cope with such stress.

As mentioned, stigmasterol also displayed a non-condensation effect. The structural difference between sitosterol and stigmasterol is only an unsaturation on C22. This has a dramatic effect on membrane fluidity as discussed in (Grosjean et al., 2015). Using model membranes and environment-sensitive probes, they showed that plant lipids promote various spatial organization of membrane and that β-sitosterol promotes *Lo* phases while stigmasterol has a low ordering effect and is correlated with low level of *Lo* phases. Plant sterols and sphingolipids form lipid rafts which are signalling platforms (Mongrand et al., 2004)(Gronnier et al., 2018). These structures can be clearly seen as lipid domains in model monolayers containing β-sitosterol and GIPC that interact with each other (Figure 7). This interaction might translate into *Lo* phases. Stigmasterol, on the other hand, tends to sequester small structures containing GIPC which might contribute to membrane fluidity.

Plants are poikilothermic and have to adapt the viscosity of their membrane to temperature changes, a process called homeoviscosity. By modulating the fluidity of their membrane to be functionally viable, plants can adapt to temperature fluctuations. For example, plants can readily convert β-sitosterol to stigmasterol by expressing the C22 desaturase CYP710 during temperature acclimation (Morikawa et al., 2006). Specific plant membrane components like β-sitosterol, stigmasterol and glucosylcerebrosides are synthesized as part of temperature adaptations to make membrane-associated biological processes possible (Beck et al., 2007). Here, we showed that GIPCs are more conducive to enable homeoviscosity. It will be interesting to further investigate how GIPCs are involved in modulating PM fluidity in thermal adaptation in synergy with other PM lipid components.

### GIPC structure in membrane organization

Recent studies provide new insight on the importance role of GIPC structure in plants through genetic approach (Fang et al., 2016; Ishikawa et al., 2018; Jiang et al., 2019; Mortimer et al., 2013). By generating mutants combined to the multidisciplinary approaches, we can uncover more about GIPC intricate structure and its biological implications. The modification of the ceramide length and hydroxylation of GIPC might alter the organization of the membrane as does sphingomyelin (SM) in animal cell, which is responsible for interdigitating between the bilayers and domain formation with cholesterol (Róg et al., 2016) (London & Brown, 2000). The closest biological molecule in terms of membrane structuring role of plant GIPC series A and B could be SM, even if the latter -absent in plant PM-is made up of a phosphocholine head group. The theoretical model of plant PM showed GIPC as the major sphingolipid in the outer leaflet, just like sphingomyelin, and inducing a lateral segregation to form *Lo* phases with phytosterols (Cacas et al., 2016)(Tjellstrom et al., 2010). Interestingly, the exact distribution of sterols in the two layers of the PM is still a matter of debate, including in animal research fields (Courtney et al., 2018). To know where sitosterol or stigmasterol are located and how they regulate the fluidity of one or both of the PM leaflets is of great interest. Unfortunately, the tools to study phytosterol distribution remain to be developed.

Plant GIPCs are structurally homologous to the animal gangliosides that are absent in plants. Gangliosides are acidic glycolipids containing sialic acid in their polar head that play an important role in immunity, signal transduction in the PM that essential for brain and retinal functions in animal cells (Sonnino & Prinetti, 2010)(Sibille et al., 2016). It is possible that polyglycosylated GIPCs have a similar role to gangliosides. Further investigation will require a better understanding of the GIPC glycosylation pattern, and the enzymes involved in GIPC biosynthesis. The present study paves the way for tackling the function of plant glycosylated sphingolipids in membrane organisation and function.

## Material and Method

### Plant Material

Cauliflower and leek were store-bought. Wild-type tobacco (cv. Bright Yellow) cell culture and rice cell culture were obtained as previously described in (Cacas et al., 2013). and (Nagano et al., 2016) respectively.

### Extraction and purification of GIPCs

The green parts of the cauliflower and leek were removed to prevent contamination by galactolipids, which are mainly present in chloroplasts. Plant material (800 g fresh weight) was blended with 5 litres of cold 0.1 N aqueous acetic acid in a chilled stainless-steel Waring Blender at low, medium and high speed for 30s each. The slurry was filtered through 16 layers of acid-washed Miracloth. The residue was re-extracted once (twice for leek) again in the same manner. The aqueous acetic acid filtrate was discarded. The residue was air-dried overnight under a fume hood and was then refluxed with 2 litres of hot 70% ethanol (0.1 N in HCl) for 20 min. The slurry was filtered hot through 16 layers of Miracloth pre-washed with acidic ethanol (pressed well to remove all liquid). This process was repeated twice more using a total of 5 litres of acidic ethanol. The combined filtrates were chilled at − 20°C for 48h. The precipitate was removed by centrifugation at 30,000 g (14000 rpm at using a Sorvall SLA-1500 rotor) at 4°C for 15 min. Sphingolipids were then extracted from the precipitates in hot isopropanol/hexane/water (55:20:25, v/v). The solution was homogenized using an Ultra-Turrax for 20 s and incubated at 60°C for 20 min. After centrifugation at 3000 g for 10 min, the supernatant was decanted to another tube and the residue extracted twice more with the hot solvent. A total of 100 ml of solvent was used at this step. The supernatants were combined and its lipid content was analysed by TLC and GC-MS to evaluate the amount of GIPC content.

Porous silica beads (Silica gel for chromatography 60 Å, 75-125 um, Acros Organics), were used throughout for packing the column chromatography. The column consists of 70 ml of silica beads, sand of Fontainebleau, followed by the sphingolipid sample dried in 20 ml of silica beads (see Figure 2). The column was washed and equilibrated with chloroform. Apolar lipids were washed with a mix of chloroform/methanol of different volume ratios of increasing polarity (4:1 then 3:1 and 2:1). The volume used was equivalent to 4-fold the volume of the column. The column was then eluted with a step gradient of chloroform:methanol:water. Solvent A was chloroform:methanol:water (59:37.5:3.5, v/v) and the solvent B chloroform:methanol:water (46:42:12, v/v). The step gradient elution started with 100% A to end with 100% B, with 10% intervals. The volume of elution corresponds to 2-fold the volume of the column. 1/100^th^ of each elution fractions were collected and dried for GC-MS and TLC analysis to test the purity of the fractions. Fractions containing the same type of GIPCs were pooled and dried. The estimated quantity of GIPC is assessed by calculating the amount of (h)VLCFA ((hydroxylated-) Very Long Chain Fatty Acid). (h)VLCFA represents 1/3 of total GIPC molecular mass.

### High performance thin-layer chromatography analysis

High-performance thin-layer chromatography (HP-TLC) plates were Silicagel 60 F254 (Merck, Rahway, NJ). HP-TLC plates were impregnated for 3 min with freshly prepared 0.2 M ammonium acetate in methanol, and further dried at 110C for 15 min. Purified lipids as well as crude extracts were chromatographed in Chloroform/Methanol/ 4N NH_4_OH (9:7:2, v/v) on. Lipids were located under UV after staining with Primuline in acetone/water 80/20.

### Carbohydrate Analysis

Samples (0.2 mg) were hydrolysed with fresh 2 M TFA at 120°C for either 1 h, 3 h, or 4 h. The supernatants were retained, dried in a vacuum concentrator, redissolved in 2 mL of water and filtered through 0.22 μm filters. Samples were analysed by HPAEC on an ICS-5000 instrument (Thermo Fisher Scientific) equipped with a CarboPac PA20 analytical anion exchange column (3 mm × 150 mm; Thermo Fisher Scientific), a PA20 guard column (3 mm × 30 mm; Thermo Fisher Scientific), a borate trap, and a pulsed amperometric detector. The column was equilibrated with 40 mM NaOH for 5 min before injection of the sample. Monosaccharides were separated using the following methods: a linear gradient from 4mM NaOH to 3 mM NaOH in the first 6 min, followed by a linear gradient of 3 mM NaOH to 1mM NaOH from 6 to 8 min. An isocratic gradient was held at 1 mM NaOH from 8 to 23 min, and then increased to 450 mM NaOH to elute the acidic sugars from 23.1 min to 45 min. Monosaccharide standards were used for quantification.

### Fatty Acid Analysis

Each sample was transmethylated at 110°C overnight in methanol containing 5% (v/v) sulfuric acid and spiked with 10 mg of heptadecanoic acid (c17:0) and 10 mg of 2-hydroxy-tetradecanoic acid (h14:0) as internal standards. After cooling, 3 mL of NaCl (2.5%, w/v) was added, and the released fatty acyl chains were extracted in hexane. Extracts were washed with 3 mL of saline solution (200 mM NaCl and 200 mM Tris, pH 8), dried under a gentle stream of nitrogen, and dissolved in 150 mL of BSTFA and trimethylchlorosilane. Free hydroxyl groups were derivatized at 110°C for 30min, surplus BSTFA-trimethylchlorosilane was evaporated under nitrogen, and samples were dissolved in hexane for analysis using GC-MS under the same conditions as described (Buré et al., 2011). Quantification of fatty acids and hydroxyl acids was based on peak areas, which were derived from total ion current, and using the respective internal standards.

### Langmuir monolayer trough

Purified GIPC-enriched fractions were used in this study. A solution at 0.4 mM in chloroform:methanol:water (30:60:8) was prepared. Sterols and PLPC were purchased from Avanti Polar Lipids. They were dissolved at 0.4 mM in chloroform:methanol (2:1). The surface pressure-area (π-A) isotherms were recorded by means of an automated Langmuir trough (KSV Minitrough [width, 75 mm; area, 24.225 mm^2^]; KSV Instruments) equipped with a platinum plate attached to a Wilhelmy-type balance. The GIPC sample was heated to 60°C for 15 min for a better solubilization. Pure solutions and lipid mixtures were spread (fixed volume of 30 μL) as tiny droplets to form a uniform monolayer on a Tris:NaCl 10:150 mM (Millipore) subphase adjusted to pH 7 with HCl. After evaporation of the solvent (15 min), monolayers were compressed at a rate of 5 mm/min and at a temperature of 22°C ± 1°C. Before each experiment, the cleanliness of the system was confirmed by checking the surface pressure over the surface compression of the pure subphase. The reproducibility of the π-A isotherms was checked by repeated recordings, and the relative SD in surface pressure and area was found to be 3% or less.

### Molecular modelling Approaches

The Hypermatrix docking procedure was used to study the interaction of GIPC with the different sterols, as already described in (Cacas et al., 2016). Briefly, one GIPC molecule is positioned and fixed for the whole calculation at the centre of the system, oriented at the hydrophobic /hydrophilic interface. The interacting molecule is also oriented at the hydrophobic/hydrophilic interface, and by rotations and translations, more than 10 million positions of the interacting molecule around the central molecule are calculated. The lowest energy matching is considered as the most stable interaction. Refer to (Cacas et al., 2016) for more details.

### Liposomes preparation (Freeze and thaw method)

The lipid solution of 1mg/ml (GIPC/PLPC or POPC or DOPC/Stigmasterol or β-sitosterol) at different molar ratio, was dried and resuspended in water. Several cycles of freeze and thaw were done with freezing occurring in liquid nitrogen for 5 min and thawing at 50°C for 15min.

### LUV preparation for DLS andζ-Potential

LUVs were prepared as described elsewhere,(Navon et al., 2017) with small modifications. Briefly, the lipid solution (GIPC/DOPC/Sterol) in 3/1 v/v THF/H_2_O methanol mixture was transferred into a round-bottom flask and the organic solvent was removed by evaporation under high vacuum pumping for 5 h, until complete evaporation of the solvent. The lipid film was then hydrated in an appropriate amount of buffer solution and subjected to 3-5 freeze thaw cycles, yielding multilamellar vesicles. The resulting suspensions (1 g L^−1^) were then successively extruded 20 times through 200 and 100 nm polycarbonate membranes using a mini-extruder (Avanti Polar Lipids).

### DLS and ζ-potential values

Dynamic Light Scattering (DLS) and ζ-Potential. DLS measurements were performed with a Malvern NanoZS instrument operating with a 2 mW HeNe laser at a wavelength of 632.8 nm and detection at an angle of 173°. All measurements were performed in a temperature-controlled chamber at 20 °C (±0.05 °C). Three measurements of 15 runs each were usually averaged. The intensity size distribution was obtained from the analysis of the correlation function using the multiple narrow mode algorithm of the Malvern DTS software. The electrophoretic mobility of the vesicles and CNCs was measured by using the same Malvern NanoZS apparatus performing at 17° from which the ζ-potential values are determined by applying the Henry equation. The ζ-potential values and the ζ-deviation were averaged over at least three measurements with at least 30 runs per measurement. They were expressed as mean ± SD (n ≥ 3).

### GUV preparation (Teflon method)

GUV were prepared as previously described by Dimova (Kubsch et al., 2017). Briefly, 50 μL of lipid mixture (1 mg mL^−1^) dissolved in organic solvent mixture were deposited on a pre-cleaned Teflon disk and the solvent was evaporated with vacuum for 2 hr. The disk was then placed in a 4 mL sealed glass vial with 200 mM sucrose and 50 mM NaCl at 60°C for 12 hours, until a cloudy deposit was formed. For microscopy observation, one volume of the vesicle suspension was mixed with 4 volumes of iso-osmolar glucose/NaCl solution for better contrast.

### Cryogenic Electronic Microscopy (Cryo-EM)

Lacey carbon formvar 300mesh copper grids were used. They were first submitted to a standard glow discharged procedure (3mbar, 3mA for 40sec). Plunge freezing was realized using the EM-GP apparatus (Leica). Four microliters of the sample was deposited on the grid and immediately blotted for 2 sec with a whatmann paper grade 5 before plunging into a liquid ethane bath cooled with liquid nitrogen (−184°C). The settings of the chamber were fixed at 70% humidity and 20°C. Total lipid concentration was 0.3 mg/ml Lipids molar ratio were as followed: POPC-^2^H_31_ / sterol (2:1), and GIPC/ POPC-^2^H_31_/Sitosterol (1:1:1). Specimens were observed at −170 °C using a cryo holder (626, Gatan, USA), with a ThermoFisher FEI Tecnai F20 electron microscope operating at 200 kV under low-dose conditions. Images were acquired with an Eagle 4k × 4k camera (ThermoFisher FEI) and processed in ImageJ. Deuterated POPC (POPC-^2^H_31_) were bought from Avanti and used as a marker for NMR measurements, GIPC were prepared from cauliflower. Sitosterol and stigmasterol were store bought from Avanti.

### Neutron Reflectivity

Neutron reflectivity experiments were performed at the ILL, on the FIGARO reflectometer (Campbell et al., 2011), on SLBs formed through vesicle fusion on silicon crystals (Montis et al., 2016; Richter at al., 2006). The crystals (dimensions l × w × h of 80 × 50 × 10 mm^3^) were polished through bath sonication in different solvents (5 min in chloroform; 5 min in acetone; 5 min in ethanol) followed by plasma cleaning. The substrates were then extensively rinsed with milliQ water and stored in milliQ water prior to use.

The specular reflectivity (R) is defined as the ratio of reflected intensity over incident intensity of a neutron beam, when the angle of reflection is equal to the angle of incidence. It is measured from a flat surface using a highly collimated neutron beam as a function of momentum transfer *Q*, where *Q* = 4*πsin0/A*, with *θ* glancing angle and *λ* wavelength. The measured reflectivity depends on the variation in the scattering length density profile, ρ(z), perpendicular to the surface. The scattering length density profile over the z-axis was modeled as a sum of discrete contributions from separate layers, each characterized by a defined scattering length density, with a gaussian roughness contribution for each interface and a solvent penetration degree. The MOTOFIT software (Nelson, 2006) which runs in the IGOR Pro environment (http://www.wavemetrics.com), was used for the analysis of the NR curves.

A multilayer model was used to analyze the reflectivity profiles of the SLBs, with fixed scattering length density values calculated for each layer: (i) a first layer of a bulk subphase of Si (ρ = 2.07 × 10^−6^Å^−2^) and a superficial layer of SiO_2_ (ρ= 3.41 × 10^−6^Å^−2^), were introduced. Their thickness and interfacial roughness were characterized in control NR measurements in D_2_O and H_2_O before vesicle injection. (ii) The polar headgroups of the SLB of the inner and outer leaflet (ρ = 1.86 × 10^−6^Å^−2^) (iii) the bilayer lipid chains (ρ = −0.30 × 10^−6^Å^−2^) (Wacklin, 2010) (iv) the sugar heads of the GIPC were represented as additional layer to the phosphate polar head group in the inner and outer leaflets (ρ = 1.9 × 10^−6^Å^−2^). (v) finally, a bulk super phase of solvent was introduced to the model.

All measurements were performed in four contrast solvents, namely H_2_O (ρ =−0.56 × 10^−6^Å^−2^), D_2_O (ρ = 6.34 × 10^−6^Å^−2^), 4MW (34% H_2_O and 66% D_2_O, ρ= 4.0 × 10^−6^Å^−2^), or SMW (62% H_2_O and 38% D_2_O,ρ= 2.07 × 10^−6^Å^−2^).

### Solid State ^2^H-NMR

Samples were prepared by co-solubilizing the appropriate amount of Bo-GIPC, POPC^2^H_31,_ sitosterol and stigmasterol in chloroform. Solvent was evaporated under a flow of nitrogen to obtain a thin lipid film, rehydrated with ultra-pure water before one-night lyophilization. The lipid powder was finally hydrated with 100μl of deuterium-depleted water (hydration of 97%). Samples were transferred into 100μl 4-mm zirconia rotors for NMR analyses. ^2^H-ssNMR experiments were performed at 76.77 MHz with a phase-cycled quadrupolar echo pulse sequence (90°x-t-90°y-t-acq) (Davis et al., 1976) and using a Bruker Avance III 500 MHz WB (11.75 T) spectrometer equipped with a solid state CPMAS 4mm H/F/X probe (IECB structural biophysics platform, Bordeaux, France). Acquisition parameters were as follows: spectral window of 500 kHz, π/2 pulse width of 3.5 μs, interpulse delays of 40 μs, recycling delay of 2s; number of scans from 1K to 6K. Spectra were processed using a Lorentzian line broadening of 300 Hz before Fourier transformation from the top of the echo. Samples were equilibrated for 20 min at a given temperature before data acquisition. All spectra were processed and analyzed using Bruker Topspin 4.0.6 software. First moments were calculated using a C^2+^ homemade routine (Buchoux S., unpublished).

### LC-MS analysis

For the analysis of sphingolipids by LC-MS/MS, lipids extracts were then incubated 1h at 50°C in 2 mL of methylamine solution (7ml methylamine 33% (w/v) in EtOH combined with 3mL of methylamine 40% (w/v) in water (Sigma Aldrich) in order to remove phospholipids. After incubation, methylamine solutions dried at 40°C under a stream of air (Markham & Jaworski, 2007). Finally, were resuspended into 100 μL of THF/MeOH/H2O (40:20:40, v/v) with 0.1% formic acid containing synthetic internal lipid standards (Cer d18:1/C17:0, GluCer d18:1/C12:0 and GM1) was added, thoroughly vortexed, incubated at 60°C for 20min, sonicated 2min and transferred into LC vials. LC-MS/MS (multiple reaction monitoring mode) analyses were performed with a model QTRAP 6500 (ABSciex) mass spectrometer coupled to a liquid chromatography system (1290 Infinity II, Agilent). Analyses were performed in the positive mode. Nitrogen was used for the curtain gas (set to 30), gas 1 (set to 30), and gas 2 (set to 10). Needle voltage was at +5500 V with needle heating at 400°C; the declustering potential was adjusted between +10 and +40 V. The collision gas was also nitrogen; collision energy varied from +15 to +60 eV on a compound-dependent basis. Reverse-phase separations were performed at 40°C on a Supercolsil ABZ+, 100×2.1 mm column and 5μm particles (Supelco). The Eluent A was THF/ACN/5 mM Ammonium formate (3/2/5 v/v/v) with 0.1% formic acid and eluent B was THF/ACN/5 mM Ammonium formate (7/2/1 v/v/v) with 0.1% formic acid. The gradient elution program for Cer and GluCer quantification was as follows: 0 to 1 min, 1%eluent B; 40 min, 80%eluent B; and 40 to 42, 80%eluent B. The gradient elution program for GIPC quantification was as follows: 0 to 1 min, 15%eluent B;31 min, 45%eluent B; 47.5 min, 70%eluent B; and 47.5 to 49, 70%eluent B. The flow rate was set at 0.2 mL/min, and 5mL sample volumes were injected. The areas of LC peaks were determined using MultiQuant software (version 3.0; ABSciex) for sphingolipids quantification, see supplemental table 1 the list of molecules Q1 ions and Q3 ions.

## Acknowledgements

We thank Jean-Paul Douliez, Catherine Sarazin and Sébastien Buchoux for critical reading and advice and Laure Beven for bright field microscopy. We thank Claire Bréhélin for the help in electronic microscopy observations. M.D. and L.L. thank the FRS-FNRS for their position as Senior Research Associates. We benefited from the facilities and expertises of the Biophysical and Structural Chemistry platform (BPCS) at IECB, CNRS UMS3033, Inserm US001, Bordeaux University http://www.iecb.ubordeaux.fr/index.php/fr/plateformestechnologiques. FS, SM, LL, LF DB are funded by the ANR PlayMobil (ANR-19-CE20-0016-02). YG, JCM were funded as was part of the DOE Joint BioEnergy Institute (http://www.jbei.org) supported by the U. S. Department of Energy, Office of Science, Office of Biological and Environmental Research, through contract DE-AC02-05CH11231 between Lawrence Berkeley National Laboratory and the U. S. Department of Energy. We thank Bordeaux-Metabolome platform for lipid analysis supported by ANR PlayMobil (grant no. ANR-19-CE20-0016-02 to S.M., L.F., F.S.-P. and Bordeaux Metabolome Facility-MetaboHUB (grant no. ANR–11–INBS–0010 to S.M. and L.F.).

## Supplementary data Figure legends

**Supplementary data 1**

High performance thin layer chromatography (HPTLC) assay of eluted fractions collected during the GIPC purification process (described in Figure 2). #2 refers to the crude extract deposited on the silica column. Fractions containing GIPC without visible contamination of sterols and phospholipids were selected and pooled to make up fraction #3.

**Supplementary data 2**

Fatty acid contents of GIPC-enriched fractions purified from cauliflower, BY-2 cell culture, leek and rice cell culture. The fatty acid content was quantified after releasing fatty acid by transmethylation of GIPC-enriched samples, followed by derivatization by BSTFA before GC-MS analysis. The most abundant pool of fatty acid are hydroxylated fatty acid >20 carbons (hVLCFA) for *Bo*-GIPC and *Nt*-GIPC enriched-samples and non-hydroxylated fatty acid >20 carbons (VLCFA) for *Ap*-GIPC and *Os*-GIPC enriched-sample. The data are means of three independent experiments. Error bars are SD.

**Supplementary data 3**

Determining glycan content by HPAE analysis of GIPC-enriched samples. A slight change in sugar amount was seen after 1h, 3h and 4h of TFA hydrolysis. (GlcA: glucuronic acid; Glc: glucose; GlcN: glucosamine; Man: mannose; Gal: galactose; Ara: arabinose; Xyl: xylose; Fuc: fucose; Rha: rhamnose; GalA: galacturonic acid).

**Supplementary data 4**

LC-MS-based sphingolidomic showing the LCB, gluCER, and GIPC content found in crude extract of cauliflower and leek, and purified Bo-GIPC and Ap-GIPC. Analysis were performed on three independent purifications and expressed as the mean relative amount in percentage of the three analysis.

**Supplementary data 5**

LC-MS-based sphingolidomic showing the different molecular species found in crude extract of cauliflower and leek, and purified Bo-GIPC and Ap-GIPC according to the LCB or (very long chain) fatty acid (VLC)FA content. Analysis were performed on triplicate of three independent purifications.The mean relative amount of each species were calculated and expressed in percentage,.

**Supplementary data 6**

Fluorescent microscopy observations of Nt-GIPC containing liposomes in water at RT, pH 7. A, Liposomes obtained after 3 cycles of freeze in liquid nitrogen and thaw (water bath at 60°C) containing (I) Nt-GIPCs 2mg/ml; (II) Nt-GIPC/ 1,2-dimyristoyl-sn-glycero-3-phosphocholine (DMPC) (4:1 mol/mol) at 2mg/ml; (III) Nt-GIPC/ DMPC (1:1 mol/mol) at 2mg/ml shows crystals; (IV) Nt-GIPC/DMPC molar ratio (1:4 mol/mol) at 2mg/ml forms liposomes of 10μm. (V) HPTLC analysis of lipid mixture confirms the presence of GIPC and DMPC in the liposome mix observed. The higher the GIPC content, the higher the occurence of crystal formation. GIPCs can form liposomes with phospholipids of short acyl chains when the latter is four-folds more abundant in the mix. Scale bar, 5μm. B, Liposomes of Bo-cauliflower GIPC/POPC/ β-sitosterol (1:1:1 mol/mol) at 1mg/ml in TBS 1X. Clusters as shown in (I) were obtained after three cycles of 20min freezing at −20°C and thawing in a water bath at 60°C. Liposomes in (II) were formed after three cycles of freezing in liquid N_2_ and 20min heating in a water bath at 60°C; no cluster was observed. The type of freeze/thaw determines the size and shape of liposomes formed. Scale bar, 5μm.

## Notes

Authors declare no conflict of interest

### Competing Interest Statement

The authors have declared no competing interest.

