## Supplementary figures and images for "Purification, characterization and influence on membrane properties of the plant-specific sphingolipids GIPC"

### supplementary data 1

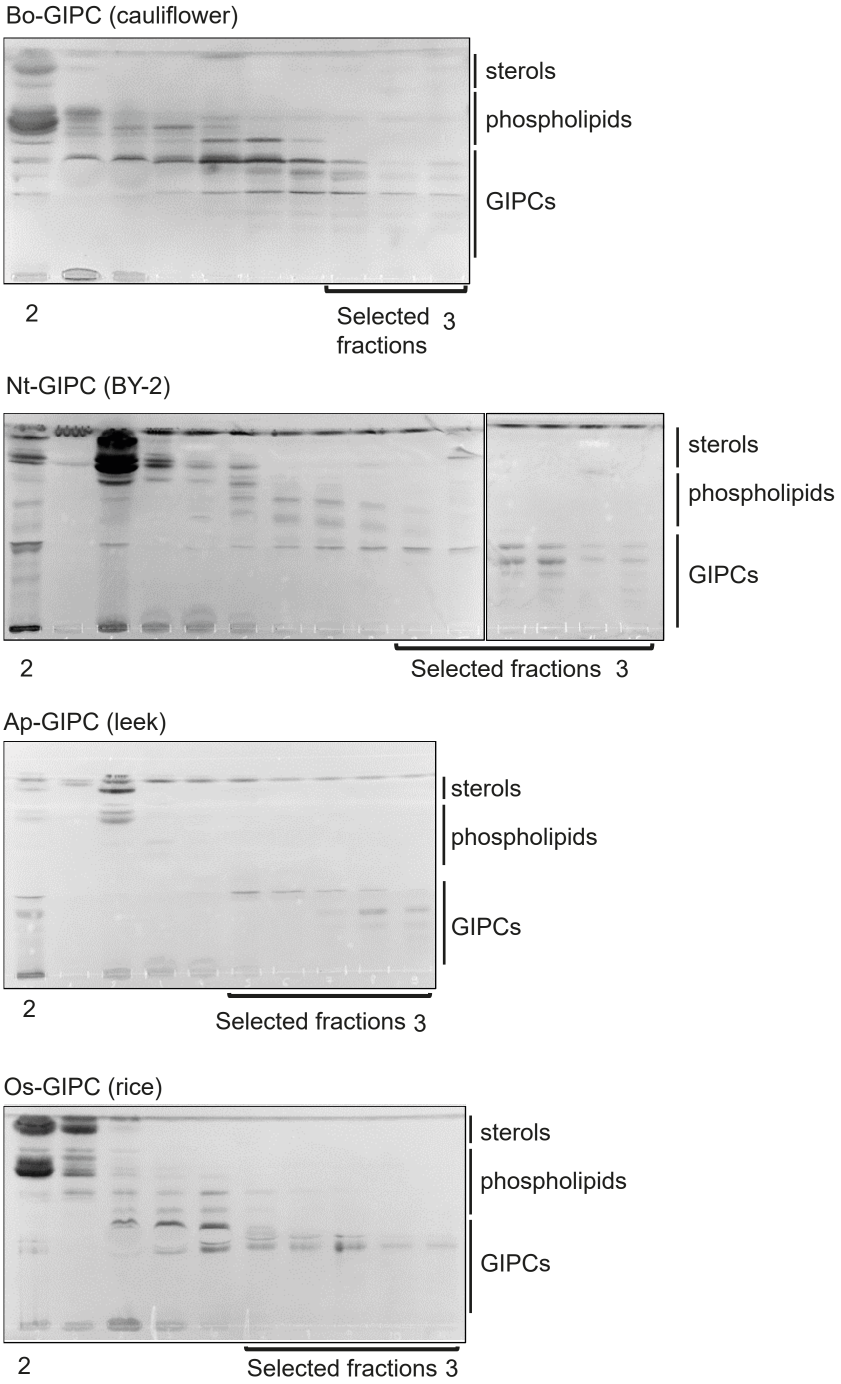

### supplementary data 2

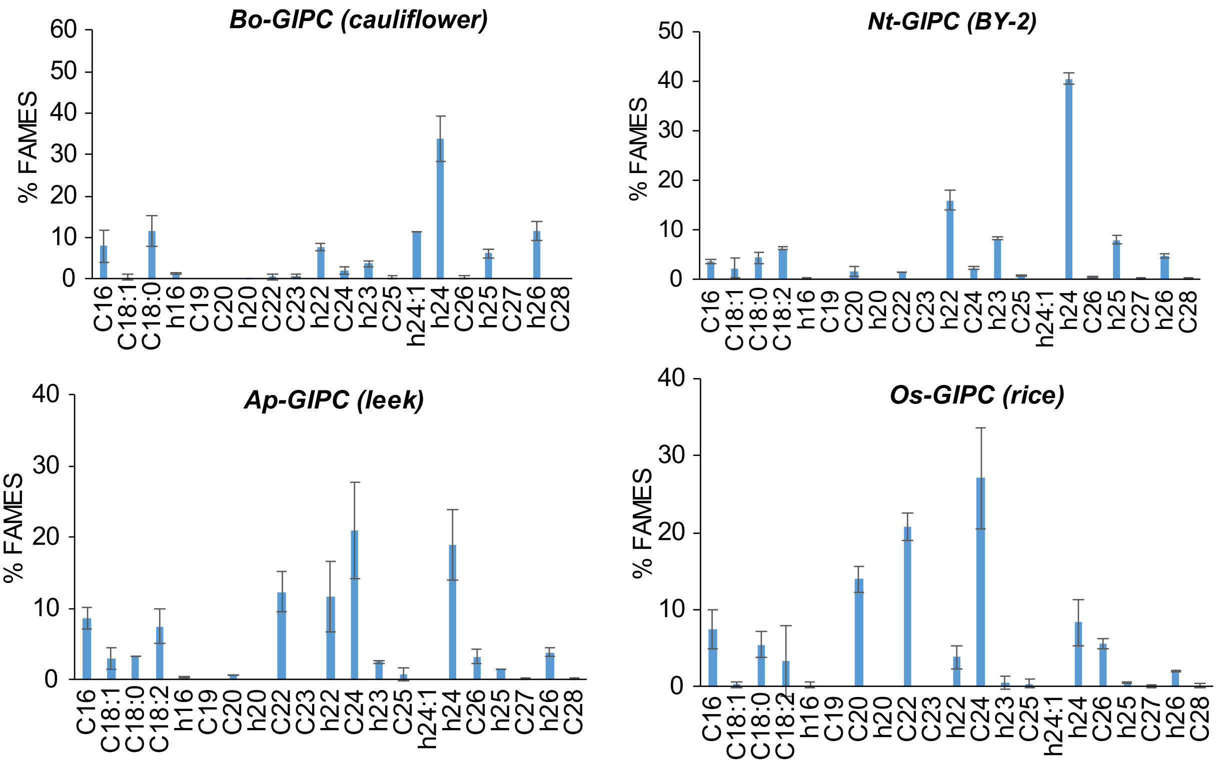

### supplementary data 3

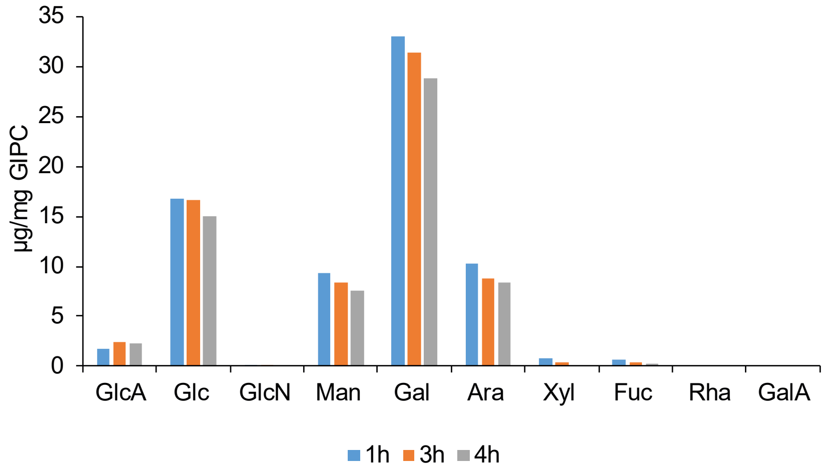

### supplementary data 5

Supplementary data 5

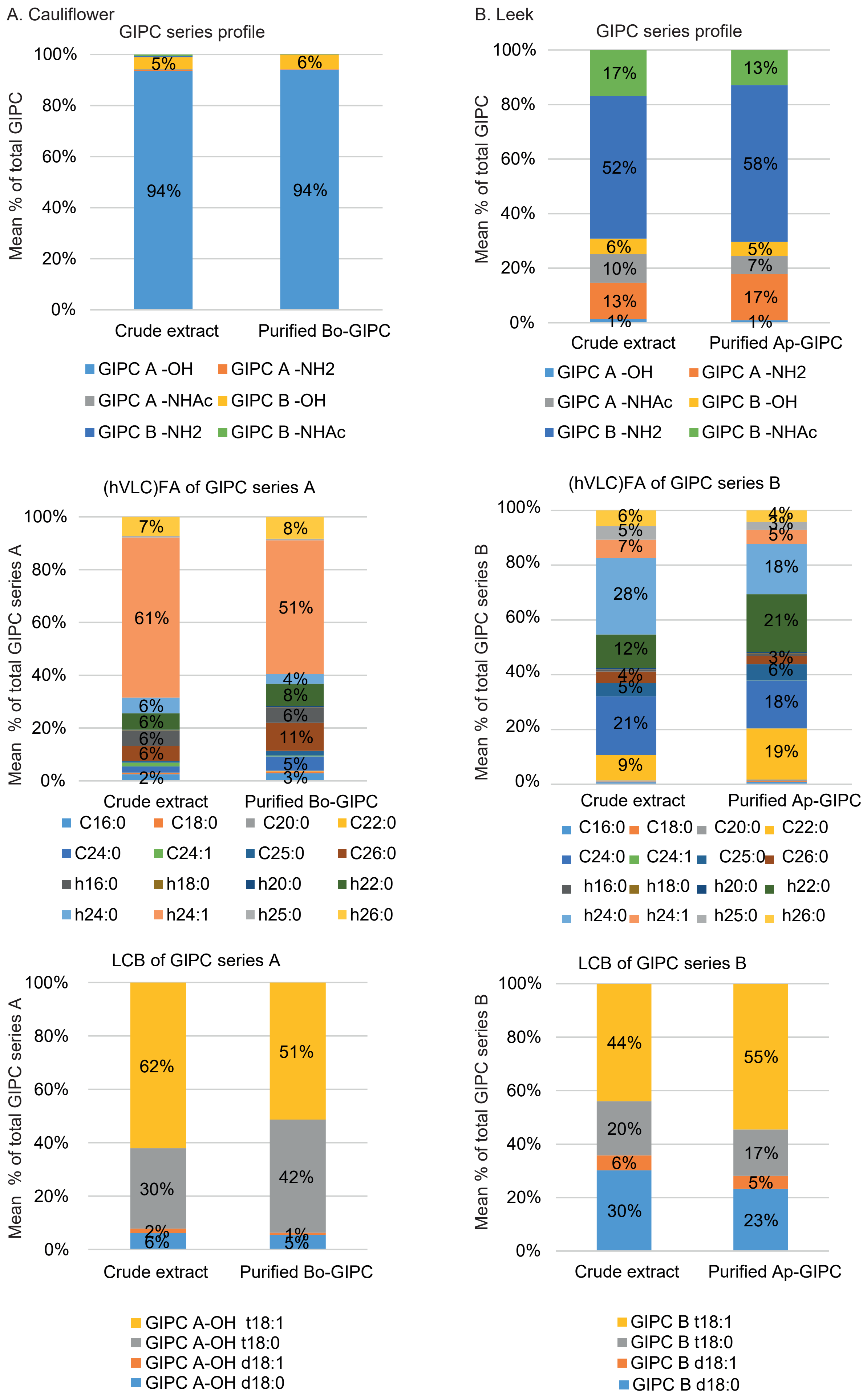

### supplementary data 6

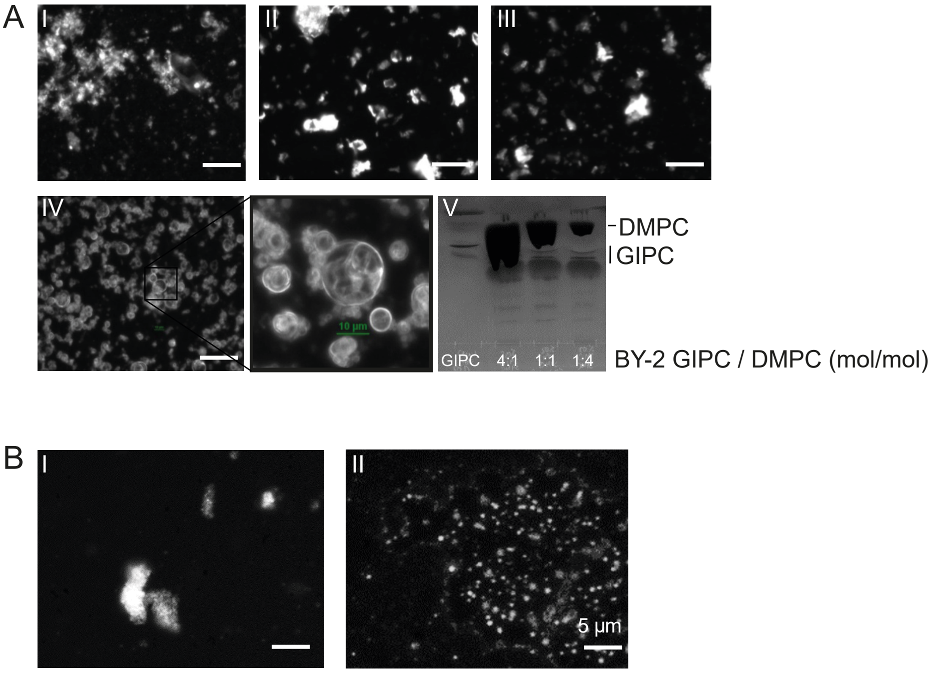
