## supplementary data 4 for "Purification, characterization and influence on membrane properties of the plant-specific sphingolipids GIPC"

Mean percentage of sphingolipid

Cauliflower

Crude extract cauliflower

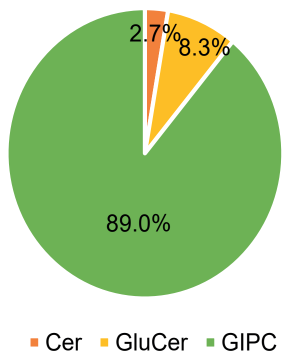


Purified Bo-GIPC

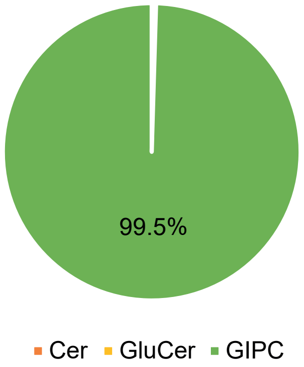


Leek

Crude extract leek

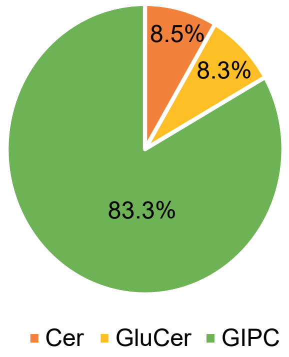


Purified Ap-GIPC

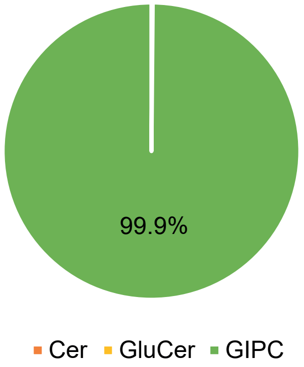
